# On the diversity of chemical power supply as a determinant of biological diversity

**DOI:** 10.1101/2020.07.05.188532

**Authors:** David Diego, Bjarte Hannisdal, Håkon Dahle

**Affiliations:** Department of Earth Science, University of Bergen, Allégaten, NO-5007 Bergen, Norway; Department of Biological Sciences, University of Bergen, Thormøhlens gate 53A, NO-5006 Bergen, Norway; K.G. Jebsen Centre for Deep Sea Research, Allégaten, NO-5007 Bergen, Norway; Computational Biology Unit, Department of Informatics, University of Bergen, N-5020 Bergen, Norway

## Abstract

Understanding how environmental factors shape biological communities is a fundamental problem in microbial ecology. Patterns of microbial diversity have been characterized across a wide range of different environmental settings, but the mechanisms generating these patterns remain poorly understood. Here, we use mathematical modelling to investigate fundamental connections between chemical power supply to a system and its biological diversity and community structure. We reveal a strong mechanistic coupling between biological diversity and the diversity of chemical power supply, but also find that different properties of power supply, such as substrate fluxes and flow and Gibbs energies of reactions, affect community structure in fundamentally different ways. Moreover, we show how simple connections between power supply and growth can give rise to complex patterns of biodiversity across physicochemical gradients, such as pH gradients. Our findings demonstrate the importance of taking into account energy fluxes in order to reveal fundamental connections between community structure and environmental variability, and to obtain a better understanding of microbial population dynamics and diversity in natural environments.

## INTRODUCTION

Numerous studies have characterized microbial diversity patterns across different environmental settings. For example, pH has been found to be a good predictor of microbial diversity in soil [1, 2] and temperature is correlated with marine planktonic bacterial richness on a global scale [3, 4], whereas salinity has been found to be correlated with microbial diversity in lake sediments [5], soil [2, 6] and estuaries [7, 8].

However, a major challenge in the field of microbial ecology is that our understanding of the underlying dynamics generating such patterns remains very limited [9–13]. Theoretical analyses of models representing the dynamics of highly idealized communities have provided useful insights into the conditions that favour co-existence of species, e.g. in terms of substrate uptake kinetics, [14–17], top down control by grazers [18, 19], and metabolic conversion of common substrates [11]. Clearly, however, the biodiversity, structure and functioning of microbial communities depend not only on species co-existence, but also on species abundances. Hence, the dynamics of abundance is of critical importance to any mechanistic account of how environmental conditions shape microbial communities and their activity.

At a fundamental level, all organisms have a demand for energy, or power, in order to grow and multiply. In principle, the available power supply should therefore represent a basic environmental constraint on the abundance of species. If this principle holds, then we would expect a strong coupling between power supply and diversity in most environments, especially under energy limited conditions. Indeed, recent gene-centric analyses of oxygen minimum zones have found that fluxes of energy seem to be robust predictors of microbial productivity and functional community structure [20, 21]. Moreover, in hydrothermal systems, the chemical energy landscapes emerging from mixing between reduced hydrothermal fluids and oxygenated cold seawater, seem to shape distributions of functional groups of bacteria and archaea [22, 23].

Environmental factors, such as pH, salinity and temperature, affect the Gibbs energies of chemical reactions, and thus modulate the chemical power supply utilized by microbial communities [24, 25]. Part of the variation in bio-diversity observed along physicochemical gradients, such as pH gradients, may therefore ultimately be linked to how those gradients affect energy landscapes.

Revealing the exact connections between chemical power supply and microbial diversity through analyses of natural environments is extremely challenging due to the high complexity and high number of unknown processes occurring in biological systems. For example, fluxes of substrate are often difficult to quantify, and extensive co-variation of variables makes it notoriously difficult to pinpoint causal effects.

An alternative to exploring natural environments is to use a theoretical modelling approach, which makes it possible to isolate the mechanistic relationship between power supply and diversity in highly idealized communities. We stress that although such models do not mimic real systems in detail, they enable us to represent basic principles in a reproducible way and to formulate testable hypotheses.

In this work, we analyse a simple population dynamics model, where growth rates are determined by maintenance powers, uptake rates of substrates, and the Gibbs energy associated with the oxidation of these substrates. We provide a thorough mathematical analysis of the relationship between biological diversity and chemical power supply in an energy limited environment. In particular, we demonstrate that complex diversity patterns along various chemical gradients can emerge from simple connections between power supply and growth. Our mathematical framework for relating chemical power supply and cellular abundances rests on fundamental thermodynamic principles.

## MATERIALS AND METHODS

### The model

We consider an idealized system, where species grow independently from each other on one limiting substrate each (Fig. 1). Hence, there is no competition for energy sources and no species-species interactions arising from food webs. We label consumers and substrates as {1, ···, *N*} such that the *i*-th consumer absorbs the *i*-th substrate. At any given instant of time *t, c_i_*(*t*) will denote the number of consumers of type *i* per unit volume present at that time. We take *c_i_* to have units of *cm*^−3^. Similarly, *s_i_*(*t*) will denote the amount of substrate of type *i* (measured in *mol*) per unit volume, so that *s_i_*(*t*) has units of mol o *cm*^−3^. Limiting substrates enter the system at fixed rates. Cellular substrate uptake rates depend on substrate concentrations in the system, and are modelled according to Michaelis-Menten kinetics as:

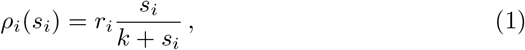

where *r_i_* (*mol · s*^−1^) denotes the maximum uptake rate and *k* (*mol · *cm**^−3^) is the half-saturation concentration^1^, that is: *ρ_i_*(*k*) = *r_i_*/2. Once absorbed by a cell, the *i*-th substrate (*S_i_*) undergoes a chemical reaction of the type *n_i_S_i_* + *a*_1_*A*_1_ + *a*_2_*A*_2_ + … + *a_n_A_n_* → *b*_1_*B*_1_ + *b*_2_*B*_2_ + …*b_m_B_m_*, with *A*_1_, ···, *A_n_* denoting any other reactants than *S_i_* and *B*_1_, ···, *B_m_* denoting products. The corresponding stoichiometric numbers are denoted by *n_i_* and *a*_1_, ···, *a_n_* and *b*_1_, ···, *b_m_*, respectively. The Gibbs energy of the chemical reaction for each mole of the substrate of type *i*, 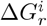, in turn depends on the substrate concentration as

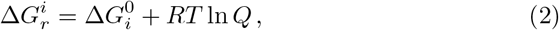

where 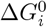 denotes the standard free Gibbs energy for the reaction, *R* is the ideal gas constant and *T* is the temperature. Moreover, *Q* denotes the reaction coefficient given by

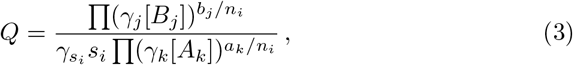

where [·] denotes the concentration (and hence *s_i_* = [*S_i_*]) and *γ*_(·)_ is the activity coefficient for the reactant/product. Letting the activity coefficients be constant for all reactants and products, and letting the concentrations be constant for all products and reactants, except for *s_i_*, we have that 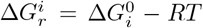 ln *s_i_* + *K_i_*, where *K_i_* is the constant 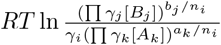. If we define an effective standard Gibbs energy as 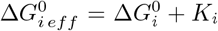, the energy available from the *i*-th reaction, which is used as an energy source by the *i*-th consumer, is

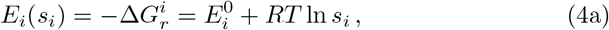

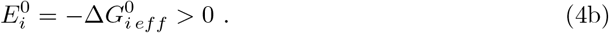

**Figure 1:**
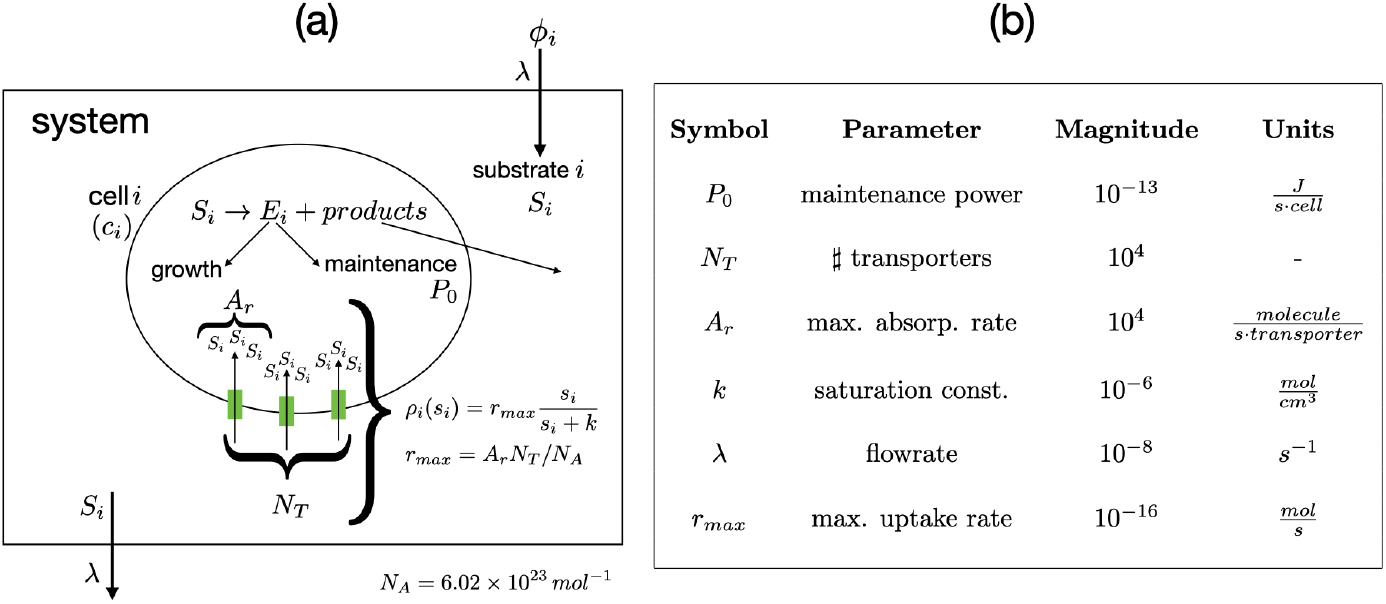
**(a) Model Schematic:** Substrates flow into and out of a system with a fixed flowrate (λ). The power supply of the *i*-th substrate is defined as the product substrate specific inflow concentration (*ϕ_i_*), the flowrate (λ) and the energy available from each mole of substrate (*E_i_*). Once the *i*-th substrate enters the system it is homogenously distributed in the system to the concentration *s_i_*. The *i*-th biological species consumes the *i*-th substrate only, and at a rate (*ρ_i_*) dependent on *s_i_*, modelled according to Michaelis-Menten kinetics in equation (1), so that the uptake of the i-th substrate by the *i*-th species is the product between *ρ_i_* and the total abundance of the *i*-th species (*c_j_*). The cell specific power supply is the product *E_i_ρ_i_*. Cellular growth rates depend on the power available for growth after a fixed amount of power has been used for maintenance (equation (6a)). The maximum uptake rate is a derived constant obtained as *r_max_* = *N_T_A_r_*/*N_A_*, with *N_A_* denoting the Avogadro number. **(b) Model parameter values**: Model constant values used in this work.

The quantity *E_i_*(*s_i_*) will be referred to as the (instantaneous) substrate-specific reaction energy and hence 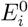 will be called the standard substrate-specific reaction energy. 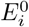 will be taken as

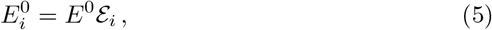

where *E*^0^ can be interpreted as the basic energy scale for the considered environment while 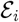 is a dimensionless factor taking into account the (possible) variability of 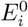 across consumers. Although values of 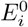 will rarely change with a common factor for all energy yielding reactions along a chemical gradient, settings with high *E*^0^ can be associated with environments where negative values of Δ*G*^0^ are typically high. Hence, *E*^0^ levels will typically be lower in anaerobic environments than in aerobic environments. However, the *E*^0^ level is also influenced by the overall chemical composition of a system so that different *E*^0^ levels may also be found along chemical gradients. As an example, consider the oxidation of lactate with sulfate as electron acceptor 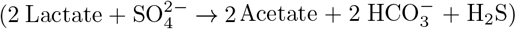, which has a standard Gibbs energy of −85.3 kJ/mol (calculated with the ‘CHNOSZ’ package in R [26]). Assuming that the activity coeffient of each reactant or product is one, *E*^0^ will shift from 236 kJ/mol in a high energy setting (acetate = 10^−3^ mM; 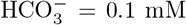; H_2_S = 0.1 mM; 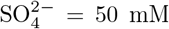) to 138 kJ/mol in a low energy setting (acetate = 50 mM; 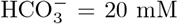; H_2_S = 10 mM; 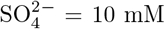). The same chemical variations would have similar effects on *E*^0^ values associated with other substrates used by sulfate reducers, such as propionate, butyrate, and ethanol.

We assume in our model that all organisms have a maintenance power demand, *P_i_* with units *J · s*^−1^. As in several previous studies [27–29], maintenance power is defined here as the power necessary to perform all cellular processes except for growth. This includes power used in spilling reactions [30–32] and power spent on ‘useful’ functions (e.g. motility). How fast the population of the i-th species grows depends on the power available for new biomass production. This power is the difference between the substrate-consumption power *E_i_*(*s_i_*) *ρ_i_*(*s_i_*), and maintenance power *P_i_*. The rate of change in substrate concentrations in the system is defined by the flow of substrate in and out of the system, as well as the rate of consumption of the substrate. This leads to the following set of ODEs:

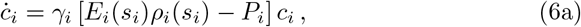

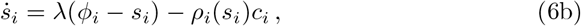

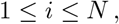

where *γ_i_*(*J*^−1^) is the biomass yield, i.e. the amount of biomass that can be built from each unit of energy. In addition, λ (*s*^−1^) is a flow rate and *φ_i_* (mol · *cm*^−3^) is the input concentration for the i-th substrate. In Fig. 1b we provide typical values for several of the model constants. Unless otherwise specified, these values are used in the simulations. In the Supplementary Information (SI) we show the main properties of the dynamics generated by the above set of differential equations. In particular, the stationary solutions correspond to

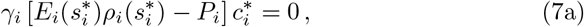

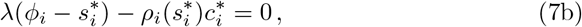

for all 1 ≤ *i* ≤ *N*. The non trivial stationary solution 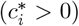 requires

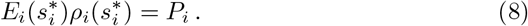

This is a transcendental equation but one may find an explicit expression for its solution in terms of the Lambert *W*-function ^2^ as (see SI for details)

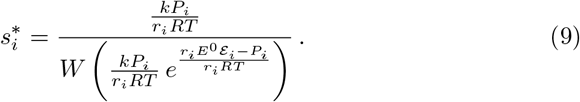

The corresponding asymptotic value for the concentration of the *i*-th consumer is then

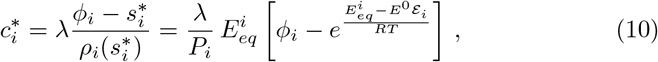

where

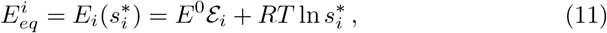

is the asymptotic value for the reaction energy corresponding to the *i*-th substrate. It can be shown that if 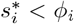, for every 1 ≤ *i* ≤ *N*, every solution to the above system with *c_i_*(0) > 0 and *s_i_*(0) > 0, for all 1 ≤ *i* ≤ *N*, verifies that 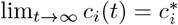 and 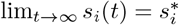, for all 1 ≤ *i* ≤ *N* (for the formal proofs, see SI).

#### Diversity

Species richness is defined as the total number of species present in an ecosystem. Species evenness, on the other hand, refers to the shape of the distribution of relative abundances of the different species. The biological diversity, i.e. the *α*-diversity, depends on both. Similarly, one can extend the concepts of richness, evenness and *α*-diversity to taxonomic groups, genes and functional groups of organisms. Here, biological *α*-diversity is defined according to the Shannon index:

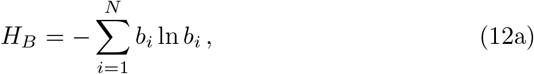

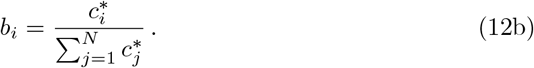

The instantaneous power supply for the *i*-th consumer is defined as

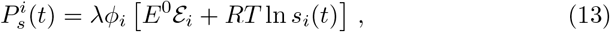

so that

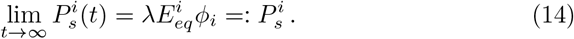

The power supply diversity is given by

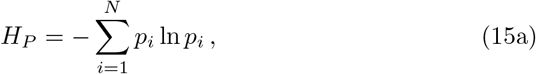

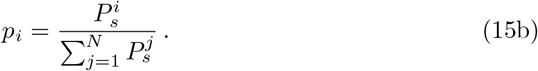

#### Variability of *H_P_* and *H_B_* across a chemical gradient

The Gibbs energy of a reaction is dependent on the activities (the product of activity coefficient and concentration) of reactants and products as in equation (2). Hence, even if the concentrations of reactants and products are kept constant, the Gibbs energy of a reaction might change due to changes in activity coefficients, which are dependent on environmental factors such as salinity. If we know how concentrations and activities vary along a physicochemical gradient, we can use equation (2) to model Gibbs energies along that gradient. We consider here the ideal case where the concentration of one compound, acting as a substrate or product in all oxidation reactions of *s_i_*, is fixed at different values across a series of independent systems. The activity coefficient of this compound is kept constant so that only variability in concentration causes variability in activity. As an example, we take such a compound to be *H*^+^. We assume the fluxes of limiting substrates to be the same for all systems. Hence, we investigate how biological diversity varies along a pH gradient. Given a substrate concentration *s_i_*, the reaction energy is thus given by

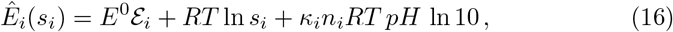

where *n_i_* is the proton stoichiometry coefficient for the reaction of the *i*-th substrate and *κ_i_* is either +1, if *H*^+^ is a product, or –1 if it is a reactant. The stationary solutions now depend explicitly on the *pH*, which entails the *pH*-dependence of both the biological and power supply diversities (see SI)

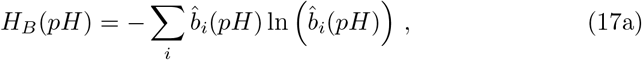

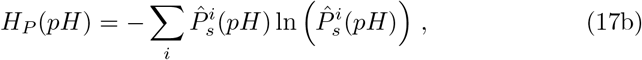

with 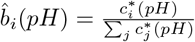 and 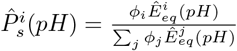.

## RESULTS

At population equilibrium, for any given species *i*, the power supply to the system 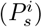 is determined by the flowrate (λ), the initial substrate concentration (*ϕ_i_*) and the reaction energy 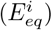 (equation (14)) whereas, in terms of the power supply 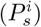, the abundance of cells for the *i*-th species 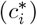 is expressed as

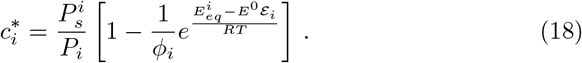

Thus, the power supply does not determine uniquely the species abundance. In order to explore the relationship between the diversity of power supply and biological diversity, we applied equations (12a) and (15a) to several combinations of *ϕ_i_*, λ, and 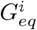 (Fig. S2).

### Relationships between biological diversity and parameters determining power supply

In this section we consider the dependence of the biological diversity on the number of consumers and on the energy they are able to extract. The number of consumers will be taken to vary on the range 10 – 1000. The variability in 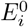, will be modelled by varying *E*^0^ on the range 10^3^ – 10^5^ *J · mol*^−1^. In addition, we will simulate several biologically relevant scenarios in terms of the availability of substrates and the efficiency of the consumers, as explained in the following.

#### Case 1: Identical power supply for all species

For reference, we first consider the trivial case where all consumers have identical traits (except for substrate specificity), and where there is no variability in input concentrations of substrates (*ϕ_i_*) or in the molar energy available from substrate oxidation (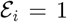 for all *i*). Clearly, in this situation all equilibrium values will be identical, in particular given by

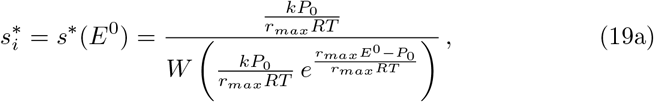

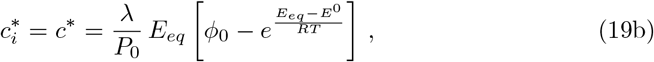

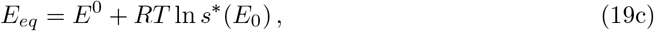

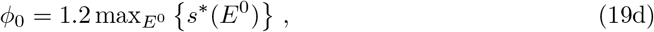

where max_E^o^_{} denotes the maximum over the range of values considered for *E*^0^. Therefore, the biological diversity (equation(12a)) will be just *H_B_* = ln *N* (*N* being the number of consumers) and hence independent of *E*^0^ (Fig. S3).

#### Case 2: Effect of variation in input concentration

In order to investigate what effect variation of input concentrations (*ϕ_i_*) has on the biological diversity, we adjust the model from case 1 so that *ϕ_i_* depends on the substrate-consumer pair as (Fig. S2a):

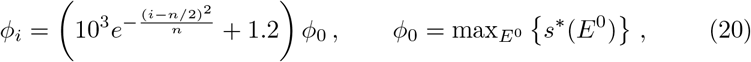

where *n* denotes the number of consumers. Due to the symmetry of these distributions around *n*/2, the relative abundances of consumers satisfy the constraint *b_i_* = *b*_*n*−*i*_. Since the asymptotic value for the concentration of substrates 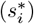 is independent of *ϕ_i_*, 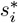 will in this case be the same for all 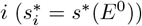 while the asymptotic values of the concentrations of consumers will in general differ as

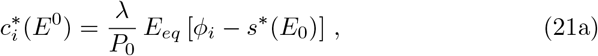

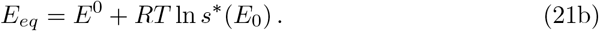

This is reflected in the decrease of the biological diversity magnitude with respect to the maximum value ln *N* (*N* being the number of consumers) (Fig. 2a). Notice that the relative abundance of each consumer is nearly independent of *E*^0^ (Fig. 3a). This can be understood as follows: From equations (22), the relative abundance for the *i*-th consumer is given by

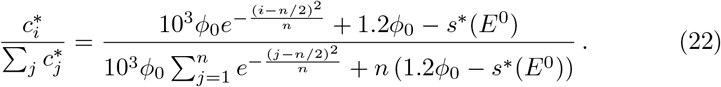

**Figure 2:**
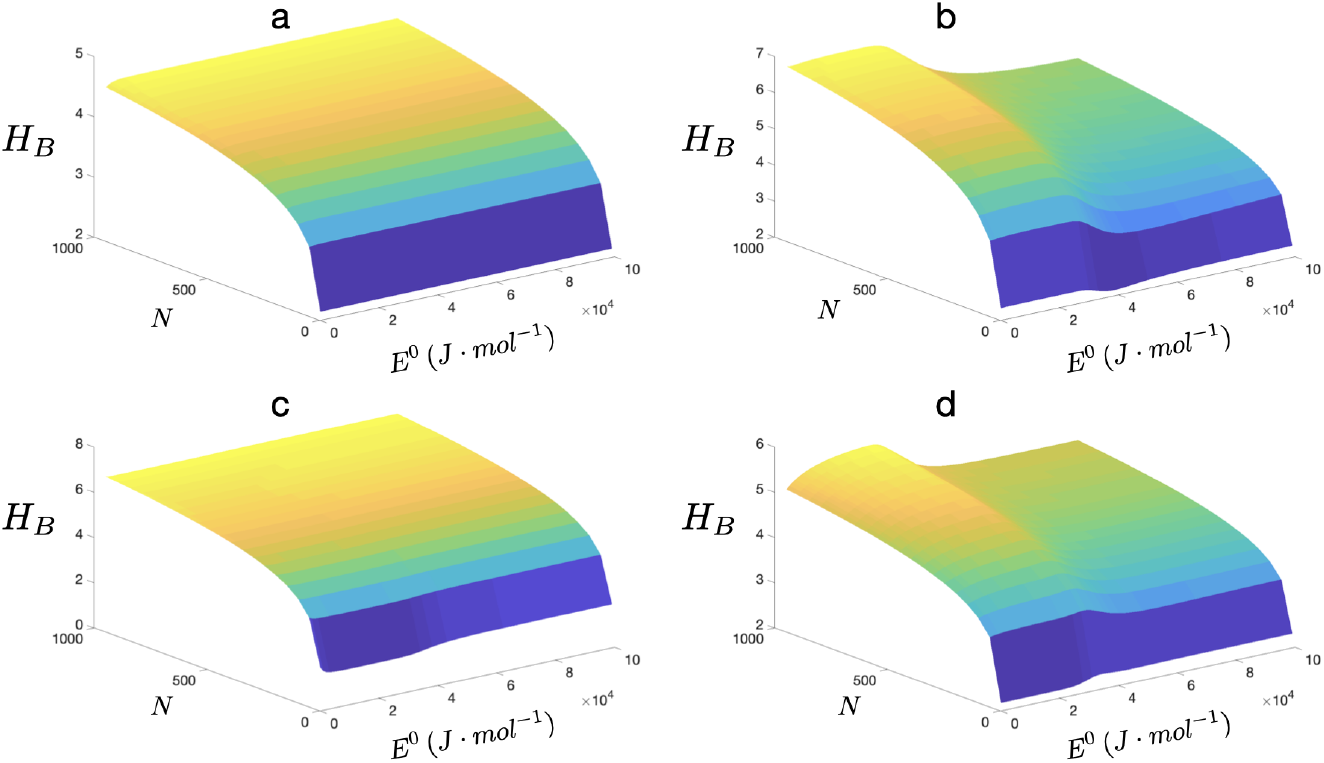
Biological diversity (*H_B_*) as a function of the number of consumers (*N*) and *E*^0^. The graphic **a** shows *H_B_* corresponding to a distribution of *ϕ_i_* given by equation (20) and model parameters as in equations (21); Graphic **b** shows *H_B_* corresponding to a distribution of 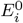 as in equation (23) and model parameters as in equations (24); Graphic **c** shows *H_B_* corresponding to distributions of *r_i_* and *P_i_* as in equations (25) and model parameters as in equations (26); The graphic **d** shows the biological diversity when all the previous distributions are considered simultaneously (model parameters as in equations (27)).

**Figure 3:**
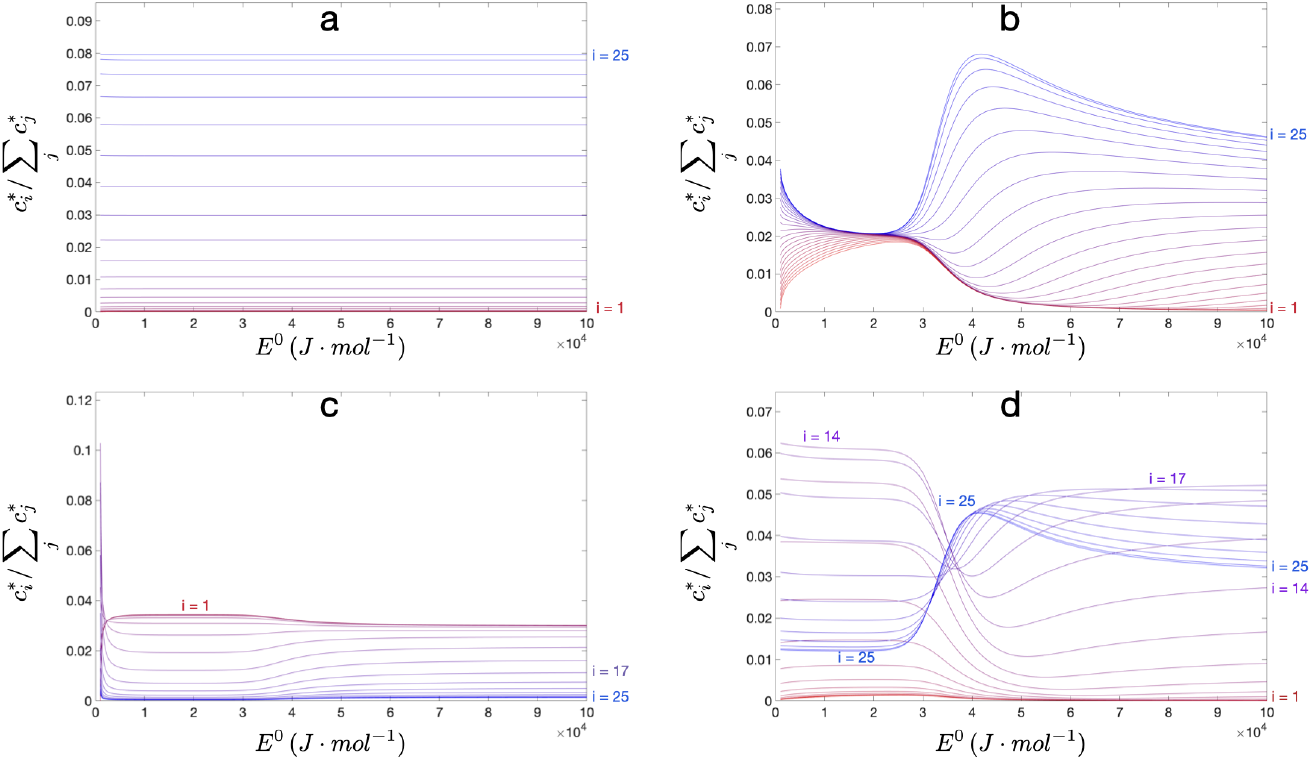
Relative abundance of cells as a function of *E*^0^ for *N* = 50 consumers. The graphics show the relative abundance corresponding to the model parameters used to produce Fig. 2a, 2b, 2c and 2d, respectively. Due to symmetry, only species labeled 1-25 are shown. The color gradient indicates species label (red - species 1; blue - species 25).

Therefore, for most of the values of *E*^0^ the dominant term in equation (22) will be 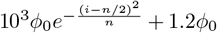 and hence the relative abundance will be nearly *E*^0^-independent. Only for low values of *E*^0^ is the term *s*^*^(*E*^0^) relevant. From equation (22) it is also clear that the most abundant consumers correspond to those having the highest supply of substrates i.e. highest *ϕ_i_* (Fig. 3a). The dependence of 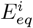 on *E*^0^ is shown in Fig. S4.

#### Case 3: Effect of variation in the energy scale across consumers

In order to investigate what effect variation in the energy level of substrate oxidation across consumers has on the biological diversity, we adjusted the model from case 1 so that *E*^0^ depends on the substrate-consumer pair as (Fig. S2b)

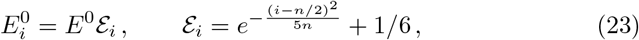

while

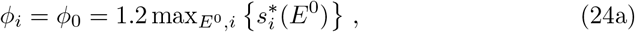

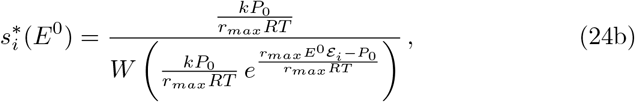

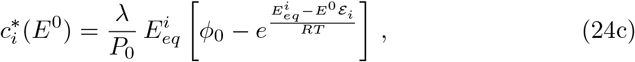

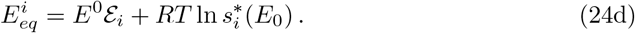

Where max_*E*^o^,*i*_ {} denotes the maximum value over the range of values for *E*^0^ and over the consumers. As expected, a gradient in the substrate reaction energy increases the sensitivity of the biological diversity on the energy scale *E*^0^ (Fig. 2b), where a clear increase of the diversity occurs for low energy scales. The relative abundance of consumers now depends on *E*^0^ in a non-trivial way (Fig. 3b). Notice that around *E*^0^ ≲ 3 × 10^4^ *J · mol*^−1^ all consumers have almost the same abundance (and hence it corresponds to the maximum value of the biological diversity in Fig. 2b). Even though the *E*^0^-dependence of the relative abundance is non-trivial, it still holds, as expected, that the most abundant consumers correspond to those with more availability of energy (Fig. 3b). The dependence of 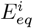 on *E*^0^ is also non-trivial in this scenario (Fig. S5).

#### Case 4: Effect of trade-off between energy acquisition efficiency and maintenance power

A biological trade-off between energy acquisition efficiency and maintenance power is arguably a key fitness trade-off in numerous habitats. For example, being motile by means of having flagella or having many highly efficient transporters will typically increase power demands (reducing the fitness), but at the same time increase the cellular power supply (increasing the fitness). Here, we model this trade-off by adjusting the case 1 model so that the distributions for the uptake rate (*r_i_*) and the maintenance power (*P_i_*) are given by (Fig. S2c and S2d)

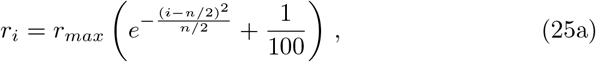

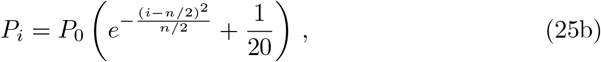

while the remaining quantities are

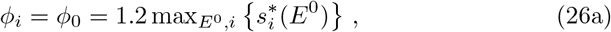

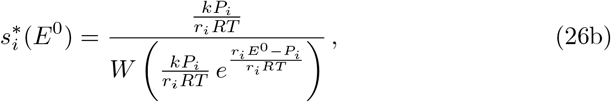

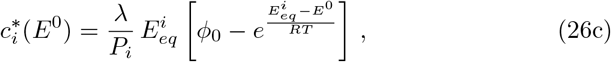

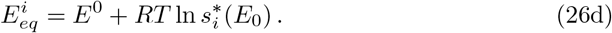

The biological diversity seems to be rather insensitive to the efficiency-cost trade-off (Fig. 2c) although the relative abundance shows a weak dependence on *E*^0^ (Fig. 3c). It is worth noting that the most abundant consumers correspond to those with low values of both uptake-rate and maintenance power (Fig. 3c), although there is little variation between species regarding the energy available from each mole of substrate 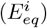 (Fig. S6).

#### Case 5: Combined effect of biological trade-off, and variability in ϕ_i_ and 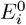

In order to investigate the combined affect of variations considered in cases 2-4, we modified the model from case 1 so that:

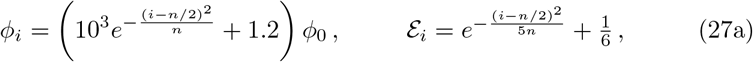

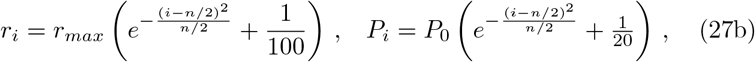

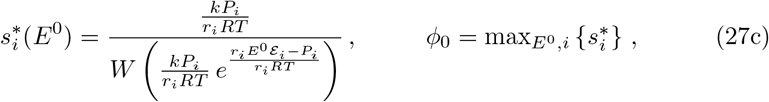

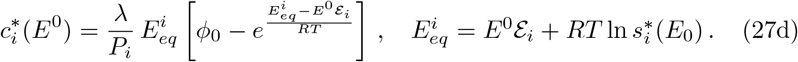

In this case, the biological diversity acquires a non-trivial *E*^0^-dependence with a clear increase towards low values of *E*^0^ (Fig. 2d). The relative abundance of consumers depends on *E*^0^ in a highly complex way (Fig. 3d). Remarkably, the particular identity of the most abundant consumer is *E*^0^-dependent (Fig. 3d). For instance, for an energy scale of *E*^0^ ~ 4 × 10^4^ *J · mol*^−1^, the most abundant consumer is the one with the highest uptake rate (*i* = 25), whereas for lower energy scales (*E*^0^ ≲ 2 × 10^4^ *J · mol*^−1^) the relative abundance of the same consumer drops from ~ 0.05 to ~ 0.01, making it one of the least abundant consumers (Fig. 3d). The complex dependence of 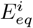 on *E*^0^ (Fig. S7) renders all values for 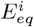 comparatively small on the range *E*^0^ ~ 2 — 4 × 10^4^ *J · mol*^−1^, while the energy availability is more markedly different across consumers for *E*^0^ ≳ 4 × 10^4^ *J · mol*^−1^, the species with the highest uptake rate (*i* = 25) being the most energetically advantaged. For *E*^0^ ≲ 2 × 10^4^ *J · mol*^−1^ however, the *i* = 25 consumer is one of the least energetically advantaged (Fig. S7). The energetic disadvantage of the *i* = 25 consumer for *E*^0^ ≲ 2 × 10^4^ *J · mol*^−1^ is clearly reflected in its low relative abundance over these energy scales (Fig. 3d). Interestingly, for *E*^0^ ≳ 5 × 10^4^ *J · mol*^−1^, the *i* = 25 consumer is not the most abundant even though it is the most energetically advantaged (i.e it has the highest *ϕ_i_* and 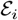 values). This asymmetric behavior across energy scales clearly emerges from the combined effect of all the above model scenarios. At a high energy scale, the percentage difference between the most energetically advantaged consumers and those with baseline values for 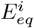, is relatively small and of little relevance. Therefore the effect of the efficiency-cost trade-off becomes more significant. This explains why, for high energy scales, the most abundant consumers correspond to those with moderate values for both *r_i_* and *P_i_* (Fig. 3d).

### Relationship between biological and power supply diversities

Here we determine how the relationship between biological diversity (*H_B_*) and power supply diversity (*H_P_*, equations (15)) is affected by distributions of *ϕ_i_*, 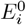, and a biological trade-off between energy acquisition efficiency and power demands. We will consider the same distributions and combinations as above (cases 2-5).

#### Case 2: Effect of variability in ϕ_i_

We find that the relation between the biological diversity (*H_B_*) and the power supply diversity (*H_P_*) is nearly linear (because *H_P_* ≃ *H_B_*) across all energy scales and number of consumers considered (Fig. 4a and Fig. S8).

**Figure 4:**
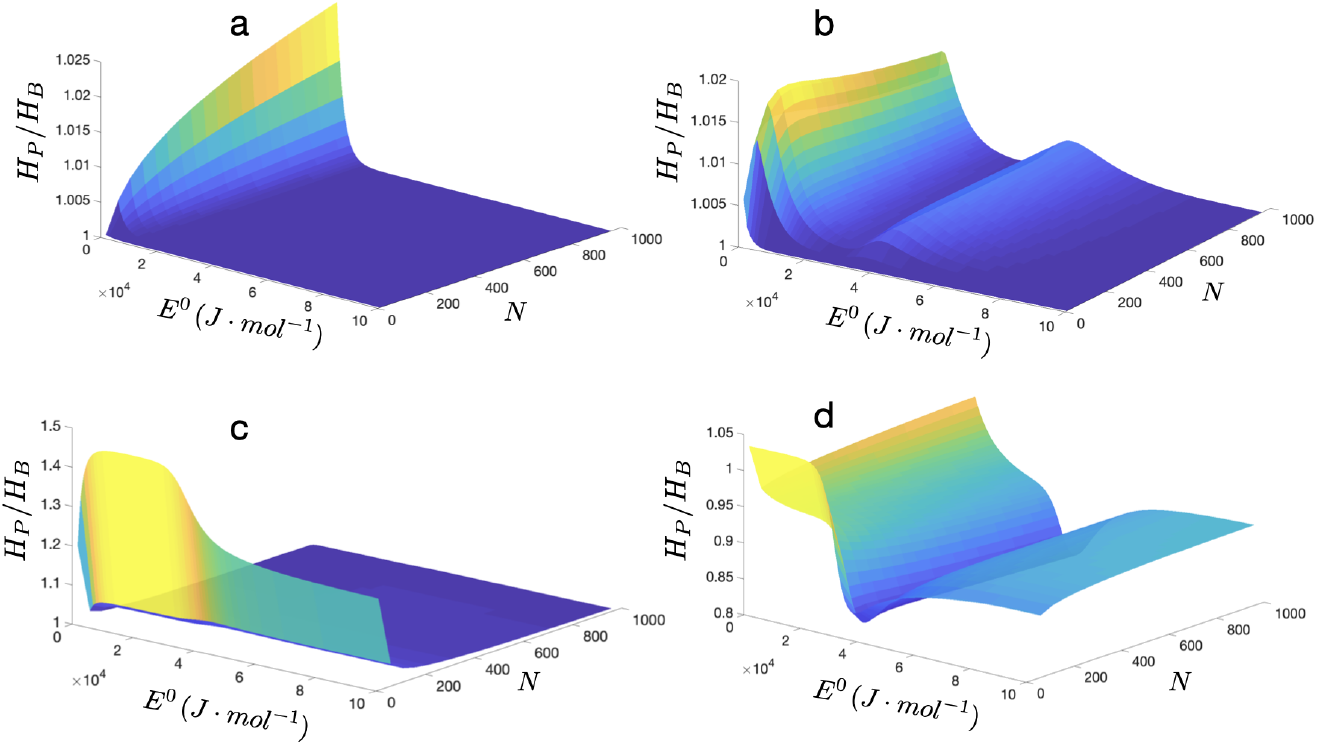
Ratio between power diversity (*H_P_*) and biological diversity (*H_B_*) as a function of the number of consumers (*N*) and the energy scale (*E*^0^). The graphics show the ratio *H_P_*/*H_B_* obtained with the model parameters used to produce Fig. 2a, 2b, 2c and 2d, respectively.

#### Case 3: Effect of variability in 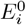

Considering instead distributions for 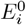 as in equation (23), we find no significant deviation from the linearity relationship *H_P_*/*H_B_* ≃ 1 over all energy scales (*E*^0^) and number of consumers (*N*) considered (Fig. 4b and Fig. S9).

#### Case 4: Effect of trade-off between energy acquisition efficiency and maintenance power

Adding a trade-off between the energy acquisition efficiency and the maintenance power increases the complexity in the relationship between HB and *H_P_*, especially for low values of the number of consumers (Fig. 4c). This implies that the relationship between *H_P_* and *H_B_* might deviate from linearity (Fig. S10). The ratio *H_P_*/*H_B_* is rather insensitive to the energy scale *E*^0^.

#### Case 5: Combined effect of biological trade-off, and variability in ϕ_i_ and 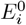

Similarly to the case 5 above, when all the above distributions are considered simultaneously, the complexity of the relation between the biological diversity and the power supply diversity increases significantly (Fig. 4d and Fig. S11). It is worth noting that each of the above considered scenarios by itself renders the power diversity always greater than the biological diversity (*H_P_*/*H_B_* ≥ 1). However, when all these scenarios are considered simultaneously, the biological diversity can become significantly greater than the power diversity (Fig. 4).

### Global scaling of the power supply

Increasing the overall power supply 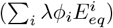 by increasing the flow of fluids into the system (i.e. increasing λ), has no effect on *H_B_*. This is evident, as λ factors out in the calculation of the relative abundance of a species (equation (12b)). However, changing the concentration of all substrates in the fluids entering the system has an effect on *H_B_*, even when the diversity of power supply *H_P_* remains unaffected. To see this, we analysed the response of the biological diversity to a global scaling of the power supply, i.e. 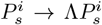, Fig. 5. In particular, under a global rescaling of the initial concentration of substrate as *ϕ_i_*(Λ) = Λ**ϕ_i_** (the Gibbs energy at equilibrium population is unaffected by such a scaling), the relative abundance of specialists (equation (12b)) is modified to

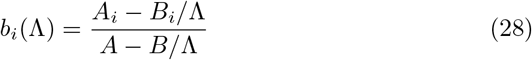

where 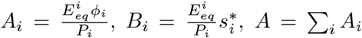 and 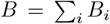. Therefore, for high enough values of the scaling factor it holds that *H_B_*(Λ) saturates to 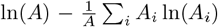 (Fig. 5). This demonstrates that changing the total power supply to the system, may or may not affect *H_B_*, depending on the exact setting of the system and whether the change in power supply is a result of changes in fluid flow or a scaling of the concentration of substrates entering the system.

**Figure 5:**
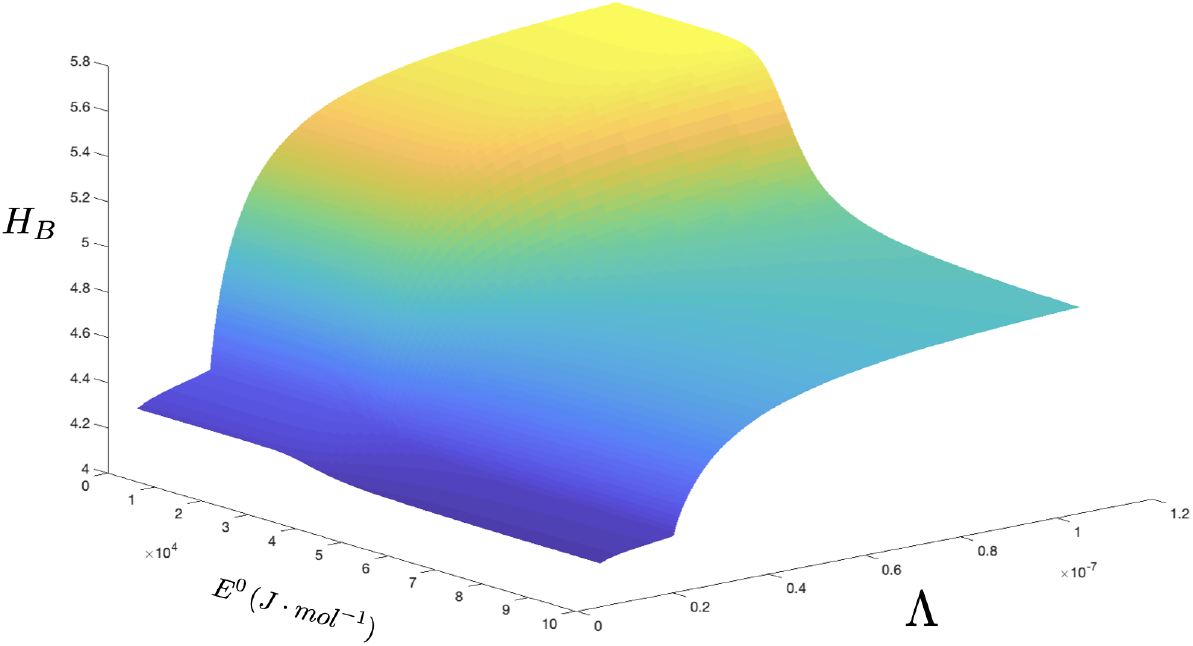
Biological diversity as a function of the energy scale (*E*^0^) and a global scaling of the input substrate concentration (i.e. *ϕ_i_* ↦ Λ*ϕ_i_*). The number of specialists is set to *n* = 500. We consider distributions for *ϕ_i_*, *r_i_* and *P_i_* given by 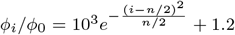, 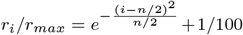 and 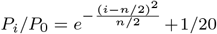. The stoichiometric coefficient is set to 5 for each substrate and the temperature is set to *T* = 300 *K*. The remaining parameters are given the values in Fig. 1b.

### Biological diversity dependence on pH

How the biological diversity changes with pH is, within our modelling framework, largely dependent on the exact chemical and biological setting of the system. For example, in a system with only two biological species where *H*^+^ is produced by one organism and consumed by the other, a decrease in pH (i.e. increase in *H*^+^) causes an increase of the power supply to the consumer of *H*^+^ whereas it makes the power supply to the producer decrease. This is easily checked using equation (16), from which we readily see that 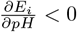 for the consumer while 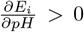 for the producer. Depending on the particular values of Δ*G*^0^ for the chemical reactions used by the two biological species, the available energies at population equilibrium 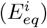 may diverge from each other (Fig. S12a) or converge to each other and cross (Fig. 6c). This reflects on the corresponding stationary concentrations of both species (Fig. 6a). The effect on the biological diversity is either a monotonic decrease (Fig. S13b) or the presence of a global maximum (Fig. 6b). In a system with many biological species, such connections may become highly complex (Fig. 6e - 6h). In particular, even when the shape of the power supply diversity is essentially the same as that for very few species (Fig. 6h and 6d), the corresponding biological diversity displays a highly complex dependence on the pH (Fig. 6f). These analyses demonstrate how *H_B_* can vary along a pH gradient, due to a thermodynamic dependency between pH and power supply. Within our modelling framework, the connections between pH and diversity will be similar between systems hosting the same biological species, but can be very different between systems hosting different biological species.

**Figure 6:**
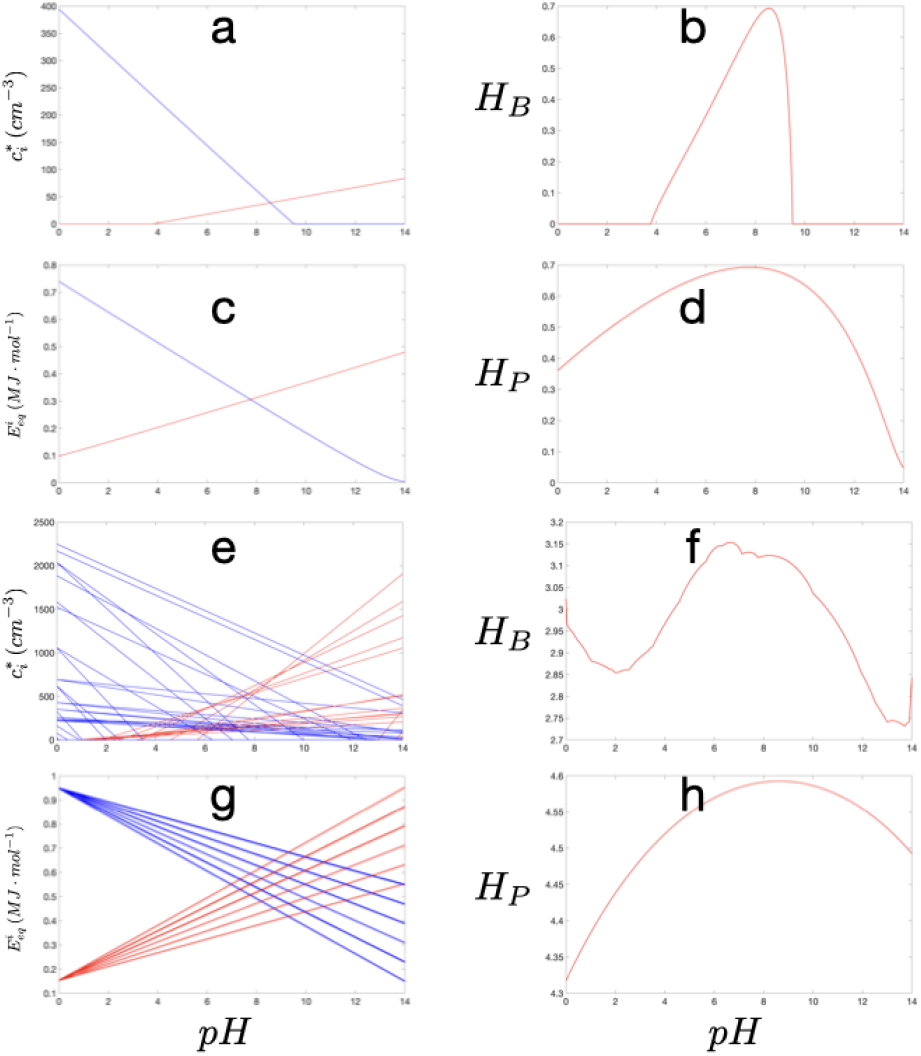
Effect of *pH* on biological and power supply diversities. The energy scale is set to *E*^0^ = 10^6^ *J · mol*^−1^ and the temperature to *T* = 300 *K*. We consider a trade-off between uptake and power maintenance as given in equations (25) while all substrates have the same input concentration given by 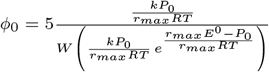, where *W*(*z*) is the Lambert *W*-function. The remaining parameters are set to the values in Fig. 1b. The graphic **a** shows the abundance of cells for the case of two specialists where one of them is an *H*^+^-producer (red line) and the other is an *H*^+^-consumer (blue line). The stoichiometric coefficients are set as 5 for the producer and 10 for the consumer; The graphic **b** shows the corresponding biological diversity; Graphic **c** shows the pH-dependence of 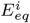 corresponding to the plot **a**. The red line shows *E_eq_* for the *H*^+^-producer and the blue line corresponds to the *H*^+^-consumer; The plot **d** shows the corresponding power supply diversity (*H_P_*); The graphics **e, f, g** and **h** show the same as **a, b, c** and **d**, respectively, but for 100 specialists. Half of them (chosen randomly) are set as *H*^+^-consumers (red lines) and the other half as *H*^+^-producers (blue lines). The stoichiometric coefficients vary between 5 and 10 and each specialist is randomly assigned a number within this interval.

## DISCUSSION

This study provides a comprehensive theoretical analysis of the coupling between fluxes of chemical energy and α-diversity. We consider a population dynamic model where growth is energy limited, which arguably is the case for most of Earth’s biosphere [33]. Our model is derived from a few fundamental principles relating chemical power supply to a system, cellular rates of substrate uptake, cellular power demands, and population size. The model assumes that biological species grow independently of each other on one limiting substrate each, hence the species richness is trivially equal to the number of limiting substrates. However, by shaping the relative abundance of species, fluxes of energy influence the biodiversity in non-intuitive ways.

The model parameters have a clear relevance to real ecosystems. For example, *λ* may describe the flow rate of substrates into and out of a fermentor or river discharge into and out of a lake; values of *ϕ_i_* describe concentrations of substrate in the inflow; *E*^0^ levels describe typical energy availability per mole of substrate oxidation under given environmental conditions, and 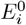 values describe cell-specific energy availability per mole of substrate. This study demonstrates that even within a simple and highly idealised model framework, complex relationships emerge between the energetic setting of a system and its biodiversity where distributions of *ϕ_i_* and 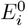, as well as *E*^0^ levels, contribute to shaping biodiversity in distinct ways. Adding a biological trade-off between energy acquisition efficiency and maintenance power increases this complexity even further.

Our numerical experiments demonstrate that a global scaling of *E*^0^ is sufficient to create changes in diversity patterns. Interestingly, *E*^0^ levels also seem to have a large impact on the identity of dominant species in models where both 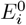 and *ϕ_i_* values vary and a trade-off between *P_i_* and *r_i_* is considered (Fig. 3d). Although values of *E*^0^ will rarely change with a common factor for all energy yielding reactions along a chemical gradient, the energy scale *E*^0^ is a potentially important parameter for understanding how environmental conditions shape the overall distribution of microbial species. Changing the power supply to a system by a scaling of *λ* has a fundamentally different effect on *H_B_* than if the same increase in power supply occurs due to a global scaling of *ϕ_i_* values - i.e. within our modelling framework, *H_B_* remains unaffected by a scaling of *λ* but responds to a scaling of *ϕ_i_* values, particularly for low *E*^0^ values Fig. 5. This finding has a clear relevance to natural systems. For example, if we want to predict the microbial diversity in an ecosystem, then the concentration of substrates in fluids flowing into the system may be a stronger predictor than the rate of fluid inflow. Note that variability in *ϕ_i_* does not affect the chemical composition of the system (except for species abundance). Consequently, environments with identical *in situ* environmental conditions may still host microbial communities with different *H_B_* due to differences in the mode of power supply.

Despite the emergent complexity of the connections between energy supply and diversity, our results suggest that the diversity of power supply (*H_P_*) may be an overall good predictor for biological diversity (*H_B_*), at least across environments with similarly shaped distributions of *ϕ_i_* and 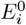 values (Fig. S8-S11). Whether or not such connections hold when more complex food webs and species interactions are considered, is clearly a topic for future research. We stress, however, that a strong correlation between *H_P_* and *H_B_* does not imply that chemical gradients shape biodiversity patterns in a simple way. Rather, as exemplified by our analysis of biodiversity along a pH gradient (Fig. 6), variations in the activity of a single chemical compound may have very different effects on *H_P_* and *H_B_* under different chemical and biological settings. Such heterogeneity in the relationship between pH and microbial *α*-diversity has been observed in different environments. In a study of 431 geographically widespread and environmentally disparate lakes, no correlation was found between *α*-diversity and pH [34]. In contrast, pH has been found to be a major driver of soil communities and is often reported to be one of the strongest predictors of *α*-diversity [1, 35]. Reported trends in the relationship between pH and microbial *α*-diversity in soil also differ. In an analysis of 300 grassland and forest soils in Germany, *α*-diversity increased with pH from pH 3 to pH 7.5, but with a plateau around pH 5 - 6 [1]. In analyses of numerous types of US soil samples, covering a pH range of 3-9, the *α*-diversity peaked at pH around 6-7. The diversity patterns observed in soils globally seem to emerge from an aggregation of multiple simpler relationships between pH and the relative abundance of individual taxonomic groups from phylum to species level [1, 36–38]. Intriguingly, this emergence of complexity from simple pH dependence of species abundance is what we find in our model (Fig. 6e,f).

Based on our modelling results, we propose three expectations that can act as working hypotheses for further inquiry:

- *E*^0^ levels and the shape of the distributions of *ϕ_i_* and 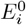 influence microbial biodiversity in different ways. *H_B_* is more sensitive to variation in the 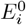 distribution than to comparable variation in the distribution of *ϕ_i_* values.
- *H_P_* is a useful predictor for *H_B_* across environments with similar *E*^0^ levels and similarly shaped distributions of *ϕ_i_* and 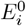.
- There is no general trend between a given chemical gradient and biodiversity, rather the relationship between them depends on the thermodynamic setting of the environment.

These expectations can be tested directly under chemostat conditions where chemical fluxes and the chemical composition of the system can be controlled, and microbial communities can be easily monitored – e.g. through 16S rRNA gene sequence analyses. In order to set up an experimental system comparable to what is modelled here, the species grown in the chemostat should have distinct substrate spectra so that each species acquires energy by the oxidation of one limiting substrate each. In principle, one could analyse diversity patterns in a system with only two species, but a higher number of species may be desirable for a more robust analysis. Estimates of maintenance power can be obtained experimentally, taking into account that maintenance power depends on environmental conditions, such as temperature [33, 39, 40].

In the field of microbial ecology, connections between environmental setting and biodiversity in natural systems have thus far mostly been explored through linear regression analyses or multivariate analyses involving directly measurable environmental parameters. Our results suggest that in order to identify driving mechanisms of biodiversity and community structure, a concerted effort should be put into assessing the role of power supply. Quantifying chemical power supply in natural environments can be challenging as it requires accurate information on chemical composition and dominant chemical fluxes in the system. Another complicating factor is that variations in the concentration of a chemical compound may have both direct and indirect effects on energy fluxes. For example, pH influences energy availability directly in energy yielding reactions where protons act as reactants or products, but also indirectly by modulating the activity coefficient or chemical speciation of numerous chemical compounds [24]. Hence, there is a need to develop improved methods for estimating energy fluxes and including such estimates in ecological studies to test model predictions.

In summary, our findings highlight the importance of taking into account energy supply and energy utilization in microbial systems in order to advance our understanding of how the fundamental laws of thermodynamics shape the biosphere.

## Acknowledgements

This work was supported by the K.G. Jebsen Center for Deep Sea Research and by a Trond Mohn Foundation Starting Grant to B.H.

## Footnotes

1 In order to reduce the multiplicity of constants, we take a common value for the half-saturation constant.

2 The Lambert *W*-function is the inverse function of *f*(*z*) = *ze^z^*.

