## Supplementary Information for "On the diversity of chemical power supply as a determinant of biological diversity"

David Diego et *al.*

### 1 Non-trivial stationary solutions

In this section we derive the expressions for the non trivial asymptotic solutions for the system of ODEs

$$\dot{c} = \gamma \left[ E(s)\rho(s) - P \right] c \,, \tag{1a}$$
$$\dot{s} = \lambda(\phi - s) - \rho(s)c \,, \tag{1b}$$

$$\rho(s) = \frac{rs}{s+k} \,,\; E(s) = E^0 + RT \ln s \,,$$

where for simplicity of the notation, we have omitted the substrate index. The solution $c^* > 0$ requires (eq. (1a))

$$E^0 + RT \ln s^* = P \frac{k + s^*}{rs^*} = \frac{P}{r} \left( \frac{k}{s^*} + 1 \right) \,, \tag{2}$$

and dividing by $RT$ and using the identity $\ln(s^*/k) = \ln s^* - \ln k$, one finds

$$\frac{E^0}{RT} + \ln k + \ln(s^*/k) = \frac{P}{r} \left( \frac{k}{s^*} + 1 \right) \,. \tag{3}$$

Defining $a = \frac{E^0}{RT} + \ln k$, $b = \frac{P}{rRT}$ and $x = \frac{s^*}{k}$, we have the equation

$$a + \ln x = b + \frac{b}{x} \,. \tag{4}$$

or equivalently

$$a - b + \ln b = \frac{1}{z} - \ln z \,, \tag{5}$$

with $z = x/b$. Taking the exponential on each side one finds

$$b\, e^{a-b} = \frac{1}{z} e^{\frac{1}{z}} \,. \tag{6}$$

Finally, in terms of $\theta = \frac{1}{z}$, one has the equality

$$\theta\, e^{\theta} = b\, e^{a-b} \,, \tag{7}$$

which has the solution

$$\theta = W \left( b\, e^{a-b} \right) \,. \tag{8}$$

where $W(y)$ is the Lambert $W$-function, which solves the equation

$$W(y)\, e^{W(y)} = y \,. \tag{9}$$

In terms of the original constants, we find

$$s^* = \frac{\frac{kP}{rRT}}{W\left(\frac{kP}{rRT}\, e^{\frac{rE^0-P}{rRT}}\right)}\,, \tag{10a}$$

$$c^* = \frac{\lambda\,(\phi - s^*)\,E(s^*)}{P}\,. \tag{10b}$$

#### 1.1 $pH$ dependence of the stationary solution

When the $pH$ is considered, the governing equation for each consumer is modified as

$$\dot{c}_i = \gamma_i \left[\hat{E}_i(s_i)\rho_i(s_i) - P_i\right] c_i\,, \tag{11}$$

with $\hat{E}_i(s_i) = \hat{E}_i^0 + RT\ln s_i$ and $\hat{E}_i^0 = E_i^0 + \kappa_i n_i RT\, pH\, \ln 10$. Therefore, the non-trivial stationary solutions will depend on the $pH$ through $\hat{E}_i^0$ as

$$s_i^*(pH) = \frac{\frac{k_iP_i}{r_iRT}}{W\left(\frac{k_iP_i}{r_iRT}\, e^{\frac{r_i\hat{E}_i^0-P_i}{r_iRT}}\right)}\,, \tag{12a}$$

$$\hat{E}^i_{eq}(pH) = \hat{E}_i^0(pH) + RT\ln\left(\hat{s}_i^*(pH)\right)\,, \tag{12b}$$

$$\hat{c}_i^*(pH) = \lambda\hat{E}^i_{eq}(pH)\frac{\phi_i - \hat{s}_i^*(pH)}{P_i}\,. \tag{12c}$$

#### 1.2 Asymptotic behavior of the Lambert $W$-function

In many instances of the numerical simulations performed in this paper the argument for the Lambert $W$-function, $z = \frac{kP}{rRT}\, e^{\frac{rE^0-P}{rRT}}$, may take exponentially large values, such as $10^{10^3}$, although this unimaginably big number is mapped by that function to a modest $\sim 10^3$. Any numerical algorithm, however, would not distinguish a number in the order of $10^{10^3}$ from $\infty$ and the evaluation would either produce a NaN (Not a Number) or a plain zero as an outcome. This problem can be avoided by noting that the argument of $W$ in our case is always of the form $z = x\,e^y$ (or equivalently $\ln z = y + \ln x$). In the following, we derive an asymptotic approximation of $W(z)$ in terms of the variable $\zeta = \ln z$.

The equation defining the $W$-function is

$$W(z)e^{W(z)} = z\,. \tag{13}$$

or equivalently

$$\ln\left(W(z)\right) + W(z) = \ln z\,. \tag{14}$$

The function $f(x) = xe^x$ has a global minimum at $x = -1$ given by $f(-1) = -\frac{1}{e}$ and in addition $\lim_{x\to+\infty} f(x) = +\infty$ while $\lim_{x\to-\infty} f(x) = 0$. Thus,

$\lim_{z\to+\infty} W(z) = +\infty$. Assume that $\ln z \gg 1$. Dividing the above expression by $\ln z$ and defining the function $\overline{W}(z) := \frac{W(z)}{\ln z}$, one finds

$$\frac{1}{\ln z}\ln\left(\overline{W}(z)\right) + \overline{W}(z) = 1 - \frac{\ln(\ln z)}{\ln z}\,. \tag{15}$$

We claim that $\overline{W}(z)$ stays bounded as $z \to \infty$. Suppose it does not. Namely, for every $M > 0$ there is $z_M > 0$ such that $\overline{W}(z_M) > M$. In particular, $W(z_M) > M\ln(z_M)$ which is possible only if $z_M \to \infty$ as $M \to \infty$. Thus, for $M$ large enough, it follows from equation (15) that

$$1 - \frac{\ln(\ln z_M)}{\ln z_M} = \frac{1}{\ln z_M}\ln\left(\overline{W}(z_M)\right) + \overline{W}(z_M) > \overline{W}(z_M) > M\,, \tag{16}$$

where we have used that $\frac{\ln(W(z_M))}{\ln z_M} > \frac{\ln M}{\ln z_M} > 0$. However, the inequality $1 - \frac{\ln(\ln z_M)}{\ln z_M} > M$ yields a contradiction as $M \to \infty$, since $\lim_{r\to\infty}\frac{\ln r}{r} = 0$.

This result tells us that there is some $K > 0$, such that $\overline{W}(z) \le K$ for all $z > 0$ , and therefore from equation (15) we find that

$$\left|\overline{W}(z) - 1 + \frac{\ln(\ln z)}{\ln z}\right| = \frac{\ln\left(\overline{W}(z)\right)}{\ln z} \le \frac{\ln K}{\ln z} \to 0\,, \tag{17}$$

as $z \to \infty$. Define $h(z) := 1 - \frac{\ln(\ln z)}{\ln z}$ and $\Omega(z) := \overline{W}(z) - h(z)$. Then, equation (15) reduces to

$$\frac{1}{\ln z}\ln\left[h(z) + \Omega(z)\right] + \Omega(z) = 0\,. \tag{18}$$

Since $\lim_{z\to\infty} h(z) = 1$ and $\lim_{z\to\infty}\Omega(z) = 0$, we may use that for $z$ large enough, $\ln(h(z) + \Omega(z)) \simeq \ln(h(z)) + \frac{1}{h(z)}\Omega(z)$, since $h(z)$ gets arbitrarily close to 1 while $\Omega(z)$ gets arbitrarily small. Hence we find the approximation

$$\frac{1}{\ln z}\left[\ln(h(z)) + \frac{1}{h(z)}\Omega(z)\right] + \Omega(z) = 0\,, \tag{19}$$

from where we find

$$\Omega(z) = \frac{W(z)}{\ln z} - h(z) \simeq -\frac{h(z)\ln(h(z))}{1 + h(z)\ln z}\,. \tag{20}$$

and therefore

$$W(z) \simeq \mathcal{W}(\ln z)\,, \tag{21a}$$

$$\mathcal{W}(\zeta) := \zeta - \ln\zeta - \{\zeta - \ln\zeta\}\,\tfrac{\ln[\zeta - \ln\zeta] - \ln\zeta}{1 + \zeta - \ln\zeta}\,. \tag{21b}$$

It turns out that this approximation is quite accurate even for not so large values of the argument. Actually, the approximation becomes accurate within 0.1% error for $z \gtrsim 100$ (Fig. S1), which corresponds to $\ln z \gtrsim 5$.

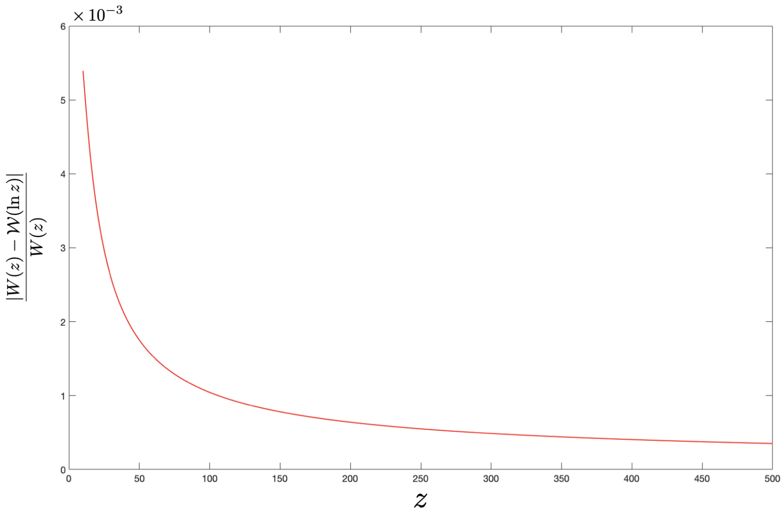

Figure S1: Relative error made by the approximation to the Lambert W-function given in equation (21). $W(z)$ represents the "exact" value (computed with the built-in MATLAB function *lambertw*) while $\mathcal{W}(\ln z)$ denotes the approximation given in equation (21).

In order to avoid the above mentioned numerical problems in the evaluation of the $W$-function, we apply the following numerical scheme:

If $y + \ln x \gtrsim 5$, then we evaluate $\mathcal{W}(y + \ln x)$.
Otherwise, we use any of the built-in implementations of the Lambert $W$-function (for instance *lambertw* in MATLAB).

#### 2 Properties of the dynamics

In this section we analyze the main properties of the dynamics generated by the differential equations

$$\dot{c}_i = \gamma_i \left[E_i(s_i)\rho_i(s_i) - P_i\right] c_i \,, \tag{22a}$$
$$\dot{s}_i = \lambda(\phi_i - s_i) - \rho_i(s_i)c_i \,, \tag{22b}$$
$$1 \leq i \leq N \,.$$

We also study the stability of the stationary solutions. The model we consider in our paper differs from the usual chemostat-like models, as those considered in [1–4], precisely on the substrate dependence of the available energy, $E_i(s) = E_i^0 + RT \ln s_i$. Accordingly, the growth term $E_i(s)\rho_i(s_i)$ for the $i$-th consumer is not simply a constant times the uptake function. Nonetheless, it turns out that with similar arguments as those used in [1–4], the same properties of the dynamics can be shown to hold. In particular the stability of the non-trivial stationary solution given in equations (10). For the sake of completeness, and even though some of the proofs proceed along rather similar lines, we find it convenient to reproduce here all the arguments leading to the final conclusion. First we note that the system of $2N$ ODEs summarized in equations (22) actually consists of $N$ independent pairs of coupled differential equations of the form

$$\dot{c} = \gamma \left[E(s)\rho(s) - P\right] c \,, \tag{23a}$$
$$\dot{s} = \lambda(\phi - s) - \rho(s)c \,. \tag{23b}$$

with $\rho(x) = \frac{rx}{k+x}$ and $E(x) = E^0 + RT \ln x$. Hence, the properties of the full dynamics may be much more easily analyzed by studying this 2-dimensional system.

We denote by $\dot{y} = V(y)$, the above 2-dimensional system, where $y = (c, s)$ and $V$ is the vector field given by $V_c(c, s) = \gamma \left[E(s)\rho(s) - P\right] c$ and $V_s(c, s) = \lambda(\phi - s) - \rho(s)c$. The stationary solutions to $\dot{y} = V(y)$ correspond to

$$\left(E(s^*)\rho(s^*) - P\right) c^* = 0 \,, \tag{24a}$$
$$\lambda(\phi - s^*) - \rho(s^*)c^* = 0 \,, \tag{24b}$$

and the non trivial stationary solution ($c^* > 0$) is given by

$$E(s^*)\rho(s^*) = P \,, \tag{25a}$$
$$c^* = \lambda \frac{\phi - s^*}{\rho(s^*)} \,. \tag{25b}$$

It is convenient to make the following definitions:

$$\mathbb{R}^2_+ := \left\{(c,s) \in \mathbb{R}^2 \,|\, c \geq 0, s > 0\right\}\,, \tag{26a}$$
$$\mathbb{K} := \left\{(c,s) \in \mathbb{R}^2_+ \,|\, c > 0,\, s < \phi\right\}\,. \tag{26b}$$

$E^* := E(s^*)$, $\rho^* := \rho(s^*)$ and $s_0 := e^{-E^0/RT}$ (i.e., $E(s_0) = 0$).

#### 2.1 General properties of the solutions

With the above definitions, we have the following results:

**Proposition 1.** *The following properties do hold.*

1. *The set $\mathbb{R}^2_+$ is forward invariant. That is: if $y(t)$ is a solution to $\dot{y} = V(y)$ with $y(0) \in \mathbb{R}^2_+$, then $y(t) \in \mathbb{R}^2_+$ for all $t \geq 0$. In particular, $s(t) > 0$ for all $t \geq 0$.*

2. *The set $\mathbb{K}$ is also forward invariant.*

*Proof.* We first show the forward invariance of $\mathbb{R}^2_+$. Let $y(t) = (c(t), s(t))$ be a solution of $\dot{y} = V(y)$ such that $c(0) = 0$ and $s(0) > 0$. The theorem of existence and uniqueness of solutions to ordinary differential equations assures that $c(t) = 0$ is the only solution to $\dot{c} = \gamma\,(E(s)\rho(s) - P)\,c$ verifying that $c(0) = 0$. In that case, the equation verified by $s(t)$ simplifies to $\dot{s} = \lambda(\phi - s)$, whose general solution is $\phi - s(t) = (\phi - c(0))e^{-\lambda t}$. In particular, $s(t) = \phi\left(1 - e^{-\lambda t}\right) + c(0)e^{-\lambda t} > 0$ for all $t \geq 0$. Suppose instead that $c(0) > 0$ and $s(0) > 0$. From the uniqueness of the solutions again, we deduce that there can not be any $t_0 > 0$, such that $c(t_0) = 0$, for then the solution would have been $c(t) = 0$ all along. Assume that $s(t_0) = 0$ for some $t_0 > 0$. More precisely, let $\tau$ be the first instant for which $s(t)$ vanishes. Then $\dot{s}(\tau) = \lambda\phi > 0$ (for $\rho(0) = 0$). The mean value theorem and the continuity of $\dot{s}$ establish that for $\epsilon > 0$, small enough, $\frac{s(\tau-\epsilon)}{-\epsilon} = \dot{s}(\tau') > 0$, for some $\tau' \in [\tau-\epsilon, \tau]$. Hence, $s(\tau-\epsilon) < 0$ which contradicts that $\tau$ is the first instant when $s(t)$ vanishes. This shows that every solution with $c(0) > 0$ and $s(0) > 0$ verifies that $c(t), s(t) > 0$ for all $t \geq 0$. In addition, the subset $\{(0,s) \,|\, s > 0\}$ is invariant and hence $\mathbb{R}^2_+$ is forward invariant.

Now let $(c(t), s(t))$ be a solution with $(c(0), s(0)) \in \mathbb{K}$ and suppose that for some $t_0 > 0$, $s(t_0) = \phi$. Then $\dot{s}(t_0) = -\rho(\phi)c(t_0) < 0$ and the same argument as before yields a contradiction. Hence $\mathbb{K}$ is forward invariant.

■

**Proposition 2.** *Every solution to $\dot{y} = V(y)$ starting in $\mathbb{R}^2_+$ is bounded.*

*Proof.* Let $y(t)$ be a solution to $\dot{y} = V(y)$ with $y(0) \in \mathbb{R}^2_+$. We claim that there are strictly positive constants $\alpha$, $\beta$, $\delta$ and $M$ such that the function $h : \mathbb{R}^2_+ \to \mathbb{R}$ given by $h(c,s) := \alpha\, c + \beta\, s + \frac{1}{2}\delta s^2$, verifies that $h(y(t)) \leq M$ for all $t \geq 0$. Since each term in $h(y(t))$ is non negative it must hold that for all $t \geq 0$, $c(t) \leq \frac{M}{\alpha}$ and $s(t) \leq \frac{M}{2\beta} + \sqrt{\frac{M}{2\delta}}$.

Now we show the claim:

Take $\eta(t) = h(y(t))$. Then $\dot{\eta}(t) = \alpha\gamma\left[E(s)\rho(s) - P\right]c + \beta\lambda(\phi - s) - \beta\rho(s)c + \delta\lambda s(\phi - s) - \delta\rho(s)sc$. We may use that for all $s > 0$, $\ln s < s$ to deduce that $E(s) < E^0 + RTs$ and therefore a first estimate is $\dot{\eta}(t) \leq \alpha\gamma E^0 c\rho(s) + \alpha\gamma RTsc\rho(s) - \alpha\gamma Pc + \beta\lambda(\phi - s) - \beta\rho(s)c + \delta\lambda s(\phi - s) - \delta\rho(s)sc$. Re-grouping the terms we find $\dot{\eta}(t) \leq -\gamma P\,\alpha c + \lambda\,(\delta\phi - \beta)\,s - \lambda\delta s^2 + \beta\lambda\phi + \left(\alpha\gamma E^0 - \beta\right)c\rho(s) + (\alpha\gamma RT - \delta)\,sc\rho(s)$. By taking $\alpha = \frac{1}{\gamma}$, $\delta = RT$ and $\beta = E^0 + \phi RT$, the function $h$ becomes $h = \frac{1}{\gamma}c + \left(E^0 + \phi RT\right)s + \frac{1}{2}RTs^2$ and the estimate for $\dot{\eta}$ reduces to $\dot{\eta}(t) \leq K - P\gamma\frac{1}{\gamma}c - \lambda E^0 s - \lambda RTs^2 - \phi RTc\rho(s) = K - P\gamma\frac{1}{\gamma}c - \frac{\lambda E^0}{E^0 + \phi RT}\left(E^0 + \phi RT\right)s - \lambda\frac{1}{2}RTs^2 - \lambda\frac{1}{2}RTs^2 - \phi RTc\rho(s) \leq$
$K - P\gamma\frac{1}{\gamma}c - \frac{\lambda E^0}{E^0 + \phi RT}\left(E^0 + \phi RT\right)s - \lambda\frac{1}{2}RTs^2$ with, $K = \lambda\phi\left(E^0 + \phi RT\right)$.
Taking $\Lambda = \min\left\{P\gamma, \frac{\lambda E^0}{E^0 + \phi RT}, \lambda\right\} > 0$, it holds that $\dot{\eta}(t) \leq K - \Lambda\eta(t)$. Thus the function $g(t) = e^{\Lambda t}\eta(t)$ verifies that $\dot{g}(t) = e^{\Lambda t}\left(\Lambda\,\eta(t) + \dot{\eta}(t)\right) \leq K\,e^{\Lambda t}$ and therefore $g(t) - g(0) \leq \frac{K}{\Lambda}\left(e^{\Lambda t} - 1\right)$ from which we deduce that $\eta(t) \leq \frac{K}{\Lambda} + \left(\eta(0) - \frac{K}{\Lambda}\right)e^{-\Lambda t}$. For $M = \max\left\{\frac{K}{\Lambda}, \eta(0)\right\}$, it therefore holds that $\eta(t) \leq M$ for all $t \geq 0$, and the claim follows.

■

**Proposition 3.** *For $P > 0$, the equality $E(s)\rho(s) = P$ has a unique solution.*

*Proof.* The function $f(x) = \frac{x}{k+x}\left(E^0 + RT\ln x\right)$ verifies that $f(x) = 0$ only when $x = s_0 = e^{-E^0/RT}$. Moreover, $f'(x) = \frac{k}{(k+x)^2}\left(E^0 + RT\ln x\right) + \frac{RT}{k+x}$ is such that $\lim_{x\to 0} f'(0) = -\infty$ and $\lim_{t\to\infty} f'(x) = +\infty$ and therefore $f'(x) = 0$ has solution. And every solution to $f'(x) = 0$ also verifies $h(x) = E^0 + RT\ln x + \frac{RT}{k}(k + x) = 0$. Its first derivative is $h'(x) = \frac{RT}{x} + \frac{RT}{k} > 0$ for all $x > 0$. Therefore $h(x) = 0$ has a unique solution over $x > 0$ and so does $f'(x) = 0$. Say that $x_0$ is the solution to $f'(x_0) = 0$. Since $\lim_{x\to 0} f(x) = 0$ and $f(s_0) = 0$ then $x_0 \in (0, s_0)$ and $f(x_0) < 0$. Moreover, since $f'(s_0) > 0$ it holds that for every $x \in (0, s_0)$, $f(x) < 0$. Suppose there is $x_1 < s_0$ such that $f(x_1) > 0$. Since $f(x_0) < 0$ there must be $x_2$ lying between $x_1$ and $x_0$ such that $f(x_2) = 0$. If $x_2 < x_0$ then we would find $x_3 < x_0$ with $f'(x_3) = 0$. If otherwise $x_2 > x_0$, there would be $x_4 > x_0$ with $f'(x_4) = 0$. In any case we contradict the fact that $f'(x) = 0$ has a unique solution. From this, it follows that the equation $E^*\rho^* = P > 0$ has a unique solution. Indeed, every value $\tilde{s}$ verifying $E(\tilde{s})\rho(\tilde{s}) = P$ must be strictly bigger than $s_0$, and if there were two such values, then we would find $\hat{s} > s_0$ such that $f'(\hat{s}) = 0$, which is a contradiction.

■

**Remark 1.** *We note that from equation (25b), $c^* > 0$ requires $s^* < \phi$ and because $P > 0$ it must hold that $s^* > s_0$.*

We now show that every orbit starting in $\mathring{\mathbb{R}}^2_+$ reaches $\mathbb{K}$ in finite time. More precisely:

**Proposition 4.** *If $s^* < \phi$, for every solution to $\dot{y} = V(y)$ with $y(0) \in \mathring{\mathbb{R}}^2_+$, there is $t_0 \geq 0$ such that $y(t_0) \in \mathbb{K}$.*

*Proof.* If $y(0) \in \mathbb{K}$, there is nothing to prove. Assume then that $y(0) \in \mathring{R}^2_+ \setminus \mathbb{K}$. This is the same as saying that $s(0) \geq \phi$. Suppose that $s(t) \geq \phi$ for all $t \geq 0$. Therefore $\dot{s}(t) \leq \lambda(\phi - s(t)) \leq 0$ for all $t \geq 0$, and since $\phi$ is a lower bound for $s(t)$ it must hold that $\lim_{t\to\infty} s(t) = \hat{s} \geq \phi$. Moreover, because $s(t) > s^*$ it holds that $E(s(t))\rho(s(t)) > E^*\rho^* = P$ for all $t \geq 0$, and hence $\dot{c} > 0$. Since the function $c(t)$ is bounded, it must follow that $\lim_{t\to\infty} c(t) = \hat{c} > c(0) > 0$, and accordingly $\lim_{t\to\infty} \dot{c}(t) = \gamma\left[E(\hat{s})\rho(\hat{s}) - P\right]\hat{c} > 0$, which is a contradiction. Thus, there must be $t_0 \geq 0$ such that $s(t_0) < \phi$ while, of course, $c(t_0) > 0$. Therefore $y(t_0) \in \mathbb{K}$, and the statement is proved.

■

To show that every orbit starting in $\mathbb{K}$ asymptotically approaches the point $(c^*, s^*)$ we still need a few more results:

**Lemma 1.** *If $s^* < \phi$, the function $f : (0, \phi) \to \mathbb{R}$ given by $f(s) := (E(s)\rho(s) - E^*\rho^*)\left[1 - \frac{\rho(s)}{\rho^*}\frac{\phi - s^*}{\phi - s}\right]$, is non positive and vanishes only at $s = s^*$.*

*Proof.* It is clear that $f(s^*) = 0$. Furthermore, since $s^* > s_0$, the equality $E(s)\rho(s) = E(s^*)\rho(s^*)$ holds only for $s = s^*$. In addition, since $E(s)R(s)$ is strictly increasing for every $s \geq s_0$ and $E(s)\rho(s) < 0$ for every $s \in (0, s_0)$, it follows that $E(s)\rho(s) > E^*\rho^*$ for every $s > s^*$ while $E(s)\rho(s) < E^*\rho^*$ for every $s \in (0, s^*)$. Therefore it holds that $\frac{E(s)\rho(s) - E^*\rho^*}{\rho(s) - \rho^*} > 0$ for every $s \neq s^*$ and accordingly, the sign of $f(s)$ equals that of $g(s) := (\rho(s) - \rho^*)\left[1 - \frac{\rho(s)}{\rho^*}\frac{\phi - s^*}{\phi - s}\right]$ for every $s \in (0, \phi) \setminus \{s^*\}$. One easily checks that for every $s \in (0, \phi) \setminus \{s^*\}$, $g(s) = -\frac{(\rho(s) - \rho^*)^2}{\rho^*}\left[1 + \frac{\rho(s)}{\phi - s}\frac{s - s^*}{\rho(s) - \rho^*}\right]$ and the mean value theorem guarantees that for some $\hat{s} \in (0, \phi)$, $\frac{\rho(s) - \rho^*}{s - s^*} = \rho'(\hat{s}) = \frac{rk}{(k + \hat{s})^2} > 0$. Hence $g(s) < 0$ over $(0, \phi) \setminus \{s^*\}$ and the lemma follows.

■

**Proposition 5.** *If $s^* < \phi$, there is a smooth function $\psi : \mathbb{K} \to \mathbb{R}$ with the properties:*

1. *$\psi \geq 0$ over $\mathbb{K}$ and vanishes only at $(c, s) = (c^*, s^*)$.*

2. *For every $p \in \left(\overline{\mathbb{K}} \setminus \mathbb{K}\right) \cap \mathbb{R}^2_+$, $\lim_{x\to p} \psi(x) = +\infty$.*

3. *$V[\psi] := V_c \frac{\partial}{\partial c}\psi + V_s \frac{\partial}{\partial s}\psi \leq 0$ over $\mathbb{K}$ and vanishes only if $s = s^*$.*

*Proof.* We claim that the function $\psi(c, s) := \frac{\phi - s^*}{\rho^*}\int_{s^*}^{s} \mathrm{d}x \frac{E(x)\rho(x) - E^*\rho^*}{\phi - x}$ $+ \frac{1}{\gamma}\left[c - c^* - c^* \ln\frac{c}{c^*}\right]$, verifies the above properties.

1. $\psi \geq 0$ over $\mathbb{K}$ and vanishes only at $(c,s) = (c^*, s^*)$.

   It clearly holds that $\psi(c^*, s^*) = 0$. In addition, the function $f : (0,\infty) \to \mathbb{R}$, given by $f(c) := c - c^* - c^* \ln \frac{c}{c^*}$, verifies that $f'(c) = 1 - \frac{c^*}{c} = 0$ only if $c = c^*$. Moreover, $f'(c) < 0$ for every $c < c^*$ and $f'(c) > 0$ for every $c > c^*$. Hence $f(c) > f(c^*) = 0$ for every $c \neq c^*$. Similarly, for the function $g : (0,\phi) \to \mathbb{R}$, given by $g(s) := \int_{s^*}^{s} \mathrm{d}x \frac{E(x)\rho(x) - E^*\rho^*}{\phi - x}$, the fundamental theorem of calculus states that $g'(s) = \frac{E(s)\rho(s) - E^*\rho^*}{\phi - s} = 0$ only if $E(s)\rho(s) = E^*\rho^*$. And because $s^* > s_0$ (for $E^* > 0$), this equality holds only when $s = s^*$. It also holds that $g'(s) > 0$ for every $s > s^*$ while $g'(s) < 0$ whenever $s < s^*$. Therefore, $g(s) > g(s^*) = 0$ for every $s \neq s^*$.

2. $\lim_{x \to p} \psi(x) = +\infty$ for every $p \in \left(\overline{\mathbb{K}} \setminus \mathbb{K}\right) \cap \mathbb{R}^2_+$.

   We note that $\left(\overline{\mathbb{K}} \setminus \mathbb{K}\right) \cap \mathbb{R}^2_+ = \left\{(c,s) \in \mathbb{R}^2_+ \mid c = 0 \vee s = \phi\right\}$. Let $p \in \left(\overline{\mathbb{K}} \setminus \mathbb{K}\right) \cap \mathbb{R}^2_+$, arbitrary. If $p_c = 0$, then $\psi(c,s) \geq c - c^* - c^* \ln \frac{c}{c^*} \to +\infty$, as $c \to 0$. Similarly, if $p_s = \phi$, then $\psi(c,s) \geq \frac{\phi - s^*}{\rho^*} \int_{s^*}^{s} \mathrm{d}x \frac{E(x)\rho(x) - E^*\rho^*}{\phi - x} \to +\infty$, as $s \to \phi$, for $E(\phi)\rho(\phi) > E^*\rho^*$.

3. $V[\psi] \leq 0$ over $\mathbb{K}$ and vanishes only if $s = s^*$.

   $V[\psi] = \frac{\phi - s^*}{\rho^*} \frac{E\rho - E^*\rho^*}{\phi - s} \left[\lambda(\phi - s) - \rho c\right] + (E\rho - P)\, c - (E\rho - P)\, c^*$. Using that $P = E^*\rho^*$ we can re-write this equality as $V[\psi] = (E\rho - E^*\rho^*) \times \left[\lambda \frac{\phi - s^*}{\rho^*} - c^*\right] + (E\rho - E^*\rho^*) \left[1 - \frac{\rho}{\rho^*} \frac{\phi - s^*}{\phi - s}\right] c$. The first term vanishes identically because $c^* = \lambda \frac{\phi - s^*}{\rho^*}$. The second term is non positive over $\mathbb{K}$ because $c > 0$ while $(E\rho - E^*\rho^*) \left[1 - \frac{\rho}{\rho^*} \frac{\phi - s^*}{\phi - s}\right] \leq 0$ for every $s \in (0,\phi)$ (lemma 1). Finally, $V[\psi] = 0$, requires $c = 0$ or $(E\rho - E^*\rho^*) \left[1 - \frac{\rho}{\rho^*} \frac{\phi - s^*}{\phi - s}\right] = 0$. The first possibility is ruled out since $(0,s) \notin \mathbb{K}$ and the second equality only holds for $s = s^*$.

■

**Proposition 6.** *For every solution to $\dot{y} = V(y)$ with $y(0) \in \mathbb{K}$, there is a compact, rectangular region $Q \subset \mathbb{K}$ such that $y(t) \in Q$ for every $t \geq 0$.*

*Proof.* We already have that $y(t) \in \mathbb{K}$ for all $t \geq 0$, for $\mathbb{K}$ is forward invariant and since $y(t)$ is bounded in $\mathbb{R}^2_+$, in particular there must be $C > 0$ such that $c(t) \leq C$ for all $t \geq 0$. In addition, the function $h = \psi \circ y$ is well defined for all $t \geq 0$, where $\psi$ is the function constructed in proposition 5. It also holds that $\dot{h}(t) = V[\psi](y(t)) \leq 0$ and hence $h(t) \leq h(0)$ for all $t \geq 0$, therefore there must be $\xi > 0$ such that $c(t) \geq \xi$ for all $t \geq 0$. Indeed, assume that such a $\xi > 0$ does not exist, that is: for every $n \in \mathbb{N}$, there is $t_n \geq 0$ such that $c(t_n) < \frac{1}{n}$. Then $h(t_n) \to +\infty$ as $n \to \infty$, which contradicts the $h$ is bounded. A similar argument shows the existence of $0 < S < \phi$ such that

$s(t) \leq S$ for all $t \geq 0$. Moreover, because $h(t)$ is decreasing ($\dot{h} \leq 0$) and bounded from below ($h(t) \geq 0$), there is $h^* \geq 0$ such that $\lim_{t\to\infty} h(t) = h^*$ and accordingly $\lim_{t\to\infty} \dot{h}(t) = 0$. From this we may deduce that there is $\sigma > 0$ such that $s(t) \geq \sigma$ for all $t \geq 0$. Indeed, if the opposite held, then for every $n \in \mathbb{N}$, there would be $t_n \geq 0$ such that $s(t_n) < \frac{1}{n}$ (or equivalently, $\lim_{n\to\infty} s(t_n) = 0$). We claim that $t_n \to \infty$, as $n \to \infty$. Indeed suppose that the sequence $\{t_n\}$ were bounded. Then we could extract a sub sequence $\{n_m\}$ with $\lim_{m\to\infty} n_m = \infty$ and such that $\lim_{m\to\infty} t_{n_m} = t_0 \geq 0$ while at the same time $s(t_0) = \lim_{m\to\infty} s(t_{n_m}) = \lim_{n\to\infty} s(t_n) = 0$, contradicting that $s = 0$ can not be reached in finite time (proposition 1). This shows the claim.

Since $\{c(t_n)\} \subset [\xi, C]$, there is a sub sequence $\{c(t_{n_m})\}$ ($\{t_{n_m}\} \to \infty$) such that $c(t_{n_m}) \to \hat{c} \in [\xi, C]$. From (the proof of) proposition 5 we know that $\dot{h}(t) = (E(s(t))\rho(s(t)) - E^*\rho^*)\left[1 - \frac{\rho(s(t))}{\rho^*}\frac{\phi - s^*}{\phi - s(t)}\right] c(t)$ and hence, $\lim_{m\to\infty} \dot{h}(t_{n_m}) = -E^*\rho^*\hat{c} < 0$ and we reach a contradiction. The proposition is proved by taking $Q := [\xi, C] \times [\sigma, S]$.

■

And finally, an easy result on $\omega$-limits of bounded curves.

**Lemma 2.** *Let $\gamma : [0, \infty) \to \mathbb{R}^n$ be a bounded curve and let $\omega_\gamma$ be its $\omega$-limit. Then for every continuous map $f : \mathbb{R}^n \to \mathbb{R}^k$, it holds that $\omega_{f\circ\gamma} = f(\omega_\gamma)$.*

*Proof.* Let $K \subset \mathbb{R}^n$, compact, such that $\gamma(t) \in K$ for all $t \geq 0$. From the continuity of $f$, it follows that $f(K)$ is also compact and it obviously holds that $f \circ \gamma(t) \in f(K)$ for all $t \geq 0$. Therefore $\emptyset \neq \omega_{f\circ\gamma} \subset f(K)$. Let $q \in f(\omega_\gamma)$, arbitrary. There is then $p \in \omega_\gamma$ such that $f(p) = q$ and also a sequence $\{t_n\} \to \infty$ such that $\{\gamma(t_n)\} \to p$. Then by the continuity of $f$ it follows that $\lim_{n\to\infty} f \circ \gamma(t_n) = f(p)$ and therefore, by definition, $q = f(p) \in \omega_{f\circ\gamma}$. Conversely, let $q \in \omega_{f\circ\gamma}$ arbitrary. There is a sequence $\{s_n\} \to \infty$ such that $\{f \circ \gamma(s_n)\} \to q$. In addition, $\{\gamma(s_n)\} \subset K$ and therefore there is a sub sequence $\{n_m\} \to \infty$ such that $\lim_{m\to\infty} \gamma(s_{n_m}) = p_* \in K$ and, by definition, $p_* \in \omega_\gamma$. Again, the continuity of $f$ implies that $q = f(p_*)$.

■

#### 2.2 Stability of the non-trivial stationary solution

In this section we show that the above non-trivial stationary solution, $(c^*, s^*)$ defined by

$$E(s^*)\rho(s^*) = P\,, \qquad c^* = \lambda\frac{\phi - s^*}{\rho(s^*)}\,,$$

is asymptotically stable with respect to $\mathring{\mathbb{R}}^2_+$, the interior of $\mathbb{R}^2_+$.

**Theorem 1.** *If $s^* < \phi$, every solution to $\dot{y} = V(y)$ with $y(0) \in \mathring{\mathbb{R}}^2_+$, verifies that* $\lim_{t\to\infty} y(t) = (c^*, s^*)$.

*Proof.* Let $x(t)$ be any solution to $\dot{x} = V(x)$ with $x(0) \in \mathring{\mathbb{R}}^2_+$. By proposition 4, there is $t_0 \geq 0$ such that $x(t_0) \in \mathbb{K}$ and hence, the map $y(t) := x(t_0 + t)$ verifies that $\dot{y} = V(y)$ and $y(0) \in \mathbb{K}$. Let $Q \subset \mathbb{K}$ be a compact region containing $y(t)$ for all $t \geq 0$. Denote by $\omega_y$, the $\omega$-limit of $y$ and by $\omega_c$ and $\omega_s$, the $\omega$-limits of $c(t) = \pi_1 \circ y(t)$ and $s(t) = \pi_2 \circ y(t)$, respectively, with $\pi_1$ and $\pi_2$ being the projections $\pi_1(c, s) = c$ and $\pi_2(c, s) = s$. By lemma 2 it holds that $\omega_c = \pi_1(\omega_y)$ and $\omega_s = \pi_2(\omega_y)$. Let $\psi : \mathbb{K} \to \mathbb{R}$ be the function constructed in proposition 5, then the function $h(t) = \psi \circ y(t)$ verifies that $\lim_{t\to\infty} h(t) = h_* \geq 0$ and hence $\lim_{t\to\infty} \dot{h}(t) = 0$. For $(\hat{c}, \hat{s}) \in \omega_y$, arbitrary, let $\{t_n\} \to \infty$ be a sequence verifying $\{y(t_n)\} \to (\hat{c}, \hat{s})$. Then, $0 = \lim_{n\to\infty} \dot{h}(t_n) = V[\psi](\hat{c}, \hat{s})$ and accordingly $\hat{s} = s^*$. Hence $\omega_s$ contains just one element $(s^*)$ and therefore $\lim_{t\to\infty} s(t) = s^*$. From this it follows that $\lim_{t\to\infty} \dot{s} = 0$ and therefore, $\lim_{t\to\infty} (c(t) - c^*) = \lim_{t\to\infty} \left\{ -\frac{\dot{s}(t)}{\rho(s(t))} + \lambda \frac{\phi - s(t)}{\rho(s(t))} - \lambda \frac{\phi - s^*}{\rho(s^*)} \right\} = 0$. Hence $\omega_c = \{c^*\}$ and therefore $\omega_y$ contains only the point $(c^*, s^*)$. This completes the proof. ∎

For completeness, we show the following result:

**Proposition 7.** *If $s^* \geq \phi$, the stationary solution $(0, \phi)$ is asymptotically stable with respect to $\mathbb{R}^2_+$.*

*Proof.* Let $x(t)$ be any solution to $\dot{x} = V(x)$ with $x(0) \in \mathbb{R}^2_+$.

If $x(0) \in \mathbb{K}$, then for all $t \geq 0$, $\dot{c}(t) < 0$, for $s(t) < \phi \leq s^*$ for all $t \geq 0$. Hence $\lim_{t\to\infty} c(t) = \hat{c} \geq 0$. However, $E(s(t))\rho(s(t)) \leq \max\{E(\phi)\rho(\phi), 0\} < P$ for all $t \geq 0$ and hence $\hat{c} = 0$. For all $\epsilon > 0$, there is $t_0 \geq 0$ such that $c(t) < \lambda \frac{\epsilon}{\rho(\phi)}$ for all $t \geq t_0$ and since $\rho(s(t)) \leq \rho(\phi)$, it holds that $\dot{s} = \lambda(\phi - s(t)) - \rho(s(t))c(t) > \lambda(\phi - \epsilon - s(t))$ for all $t \geq t_0$. The inequality $\dot{s}(t) > \lambda(\phi - \epsilon) - \lambda s(t)$ can be integrated to $s(t) > \phi - \epsilon + [s(t_0) - (\phi - \epsilon)]\, e^{\lambda(t_0 - t)}$ for all $t \geq t_0$. If $s(t_0) \geq \phi - \epsilon$ we already have that $s(t) > \phi - \epsilon$ for all $t \geq t_0$. If otherwise $s(t_0) < \phi - \epsilon$, then there is $t_1 > t_0$ such that $[\phi - \epsilon - s(t_0)]\, e^{\lambda(t_0 - t)} < \epsilon$ for all $t \geq t_1$, and therefore $s(t) > \phi - \epsilon - [\phi - \epsilon - s(t_0)]\, e^{\lambda(t_0 - t)} > \phi - 2\epsilon$ for all $t \geq t_1$. Since $\epsilon$ can be taken arbitrarily small, it follows that $s(t) \to \phi$. If otherwise $x(0) \in \mathbb{R}^2_+ \setminus \mathbb{K}$ then either $c(0) = 0$ or $s(0) \geq \phi$ and $c(0) > 0$. In the first case, $c(t) = 0$ is the only possible solution with that initial value and hence $\dot{s} = \lambda(\phi - s)$ which implies $s(t) \to \phi$. Suppose otherwise, that $s(0) \geq \phi$ and $c(0) > 0$. It clearly holds that either $s(t) \geq \phi$ for all $t \geq 0$ or there is $t_2 > 0$ such that $s(t_2) < \phi$. In the second possibility, defining $y(t) := x(t_2 + t)$ and using the same argument as before we conclude that $x(t) \to (0, \phi)$. If instead $s(t) \geq \phi$ for all $t \geq 0$, then $\dot{s} = \lambda(\phi - s(t)) - \rho(s(t))c(t) \leq 0$ for all $t \geq 0$ and therefore $\lim_{t\to\infty} s(t) = \hat{s} \geq \phi$ and also $\lim_{t\to\infty} \dot{s}(t) = 0$. Denote by $\omega_x$, the $\omega$-limit of $x(t)$ (since $x(t)$ is bounded, $\emptyset \neq \omega_x \subset \overline{\mathbb{R}^2_+}$) and let $p \in \omega_x$ arbitrary. From what we just saw, $p = (\hat{c}, \hat{s})$, with $\hat{s} \geq \phi$ and of course $\hat{c} \geq 0$. Then for some sequence $\{t_n\} \to \infty$ it holds that $0 = \lim_{n\to\infty} \dot{s}(t_n) = \lambda(\phi - \hat{s}) - \rho(\hat{s})\hat{c} \leq \lambda(\phi - \hat{s})$ and thus $\hat{s} \leq \phi$ which requires $\hat{s} = \phi$. Accordingly $\lim_{t\to\infty} s(t) = \phi$. If $s^* > \phi$, there is $t_3 \geq 0$ such that $s(t) < s^*$ for all $t \geq t_3$ and therefore $\dot{c}(t) \leq 0$ for all $t \geq t_3$

(for $E(s(t))\rho(s(t)) < E^*\rho^* = P$ for all $t \geq t_3$). Therefore $\lim_{t\to\infty} c(t) = \hat{c} \geq 0$ and $\lim_{t\to\infty} \dot{c}(t) = \gamma\,[E(\phi)\rho(\phi) - P]\,\hat{c} = 0$. If $\hat{c} > 0$, then $E(\phi)\rho(\phi) = P$, which is a contradiction. If otherwise $s^* = \phi$, since we are assuming that $s(t) \geq \phi$, for all $t \geq 0$, it still holds that $\lim_{t\to\infty} s(t) = \phi = s^*$ and accordingly $E(s(t))\rho(s(t)) \geq P$ for all $t \geq 0$ and so that $\dot{c}(t) \geq 0$. From the boundedness of $c(t)$ it must follow that $\lim_{t\to\infty} c(t) = \hat{c} \geq c(0) > 0$, but this yields the contradiction $\lim_{t\to\infty} \dot{s}(t) = -\rho(\phi)\hat{c} < 0$. From this contradiction, we deduce that if $s^* = \phi$ and $s(0) \geq \phi$ and $c(0) > 0$, then there must be $t' \geq 0$ such that $s(t') < \phi$. Therefore $x(t') \in \mathbb{K}$ and $\lim_{t\to\infty} x(t) = (0, \phi)$ also holds. ∎

### 3 Supplementary figures

#### 3.1 Substrate-specific model parameters

We consider distributions of input substrate concentration ($\phi_i$), substrate-specific reaction-energy ($E_i^0$), uptake rate ($r_i$) and maintenance power ($P_i$) given by

$$\phi_i = \left(10^3 e^{-\frac{(i-n/2)^2}{\sigma n}} + 1.2\right)\phi_0 \,, \quad E_i^0 = \left(e^{-\frac{(i-n/2)^2}{5\sigma n}} + \tfrac{1}{6}\right)E^0 \,, \tag{27a}$$

$$r_i = r_{max}\left(e^{-\frac{(i-n/2)^2}{\sigma n/2}} + \frac{1}{100}\right) \,, \quad P_i = P_0\left(e^{-\frac{(i-n/2)^2}{\sigma n/2}} + \tfrac{1}{20}\right) \,, \tag{27b}$$

where $\sigma$ controls the distribution width.

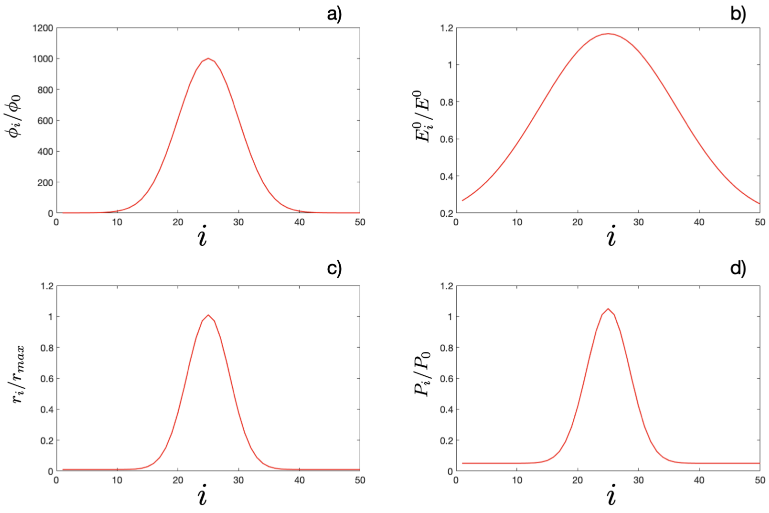

Figure S2: Typical distributions of input substrate $\phi_i$ (graphic a), standard substrate reaction energy $E_i^0$ (graphic b), uptake rate $r_i$ (graphic c) and maintenance power $P_i$ (graphic d) used for the numerical simulations presented in this work (see main text).

324 ## 3.2 Biological diversity as a function of the energy scale
325 $E^0$ and the number of consumers $N$

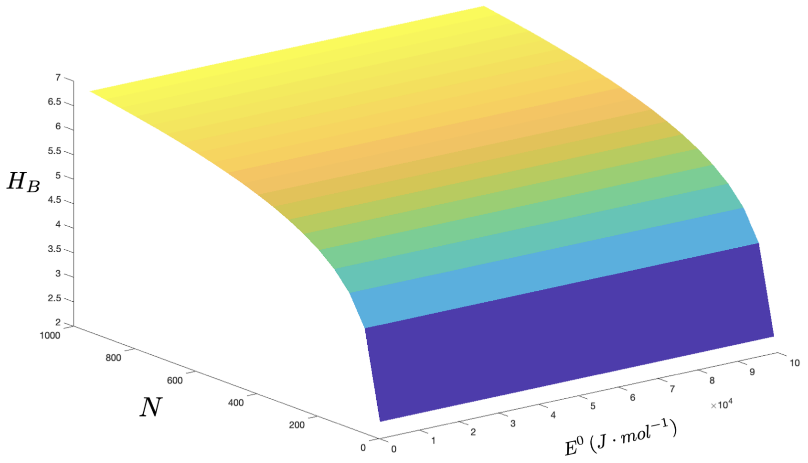

Figure S3: Biological diversity ($H_B$) as a function of the number of consumers ($N$) and the energy scale ($E^0$). All model parameters are set as common to all consumers.

#### 3.3 Available energy at population equilibrium

We consider the dependence of the available energy at population equilibrium ($E^i_{eq} = E^0_i + RT \ln s^*_i(E_0)$) as a function of the energy scale ($E^0$) for distributions of input substrate concentration ($\phi_i$), substrate-specific reaction-energy ($E^0_i$), uptake rate ($r_i$) and maintenance power ($P_i$) as in Fig. S2.

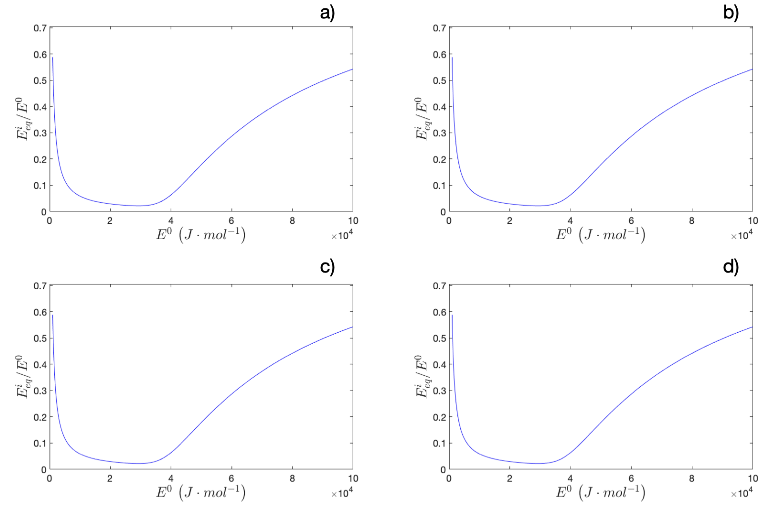

Figure S4: Available energy at population equilibrium ($E^i_{eq}/E^0$) as a function of $E^0$ for $N = 50$ consumers and $\phi_i$ distributions as in Fig. S2a for several widths: a) $\sigma = 0.5$ ; b) $\sigma = 1$ ; c) $\sigma = 2$ ; d) $\sigma = 4$. The remaining model parameters are common to all specialists.

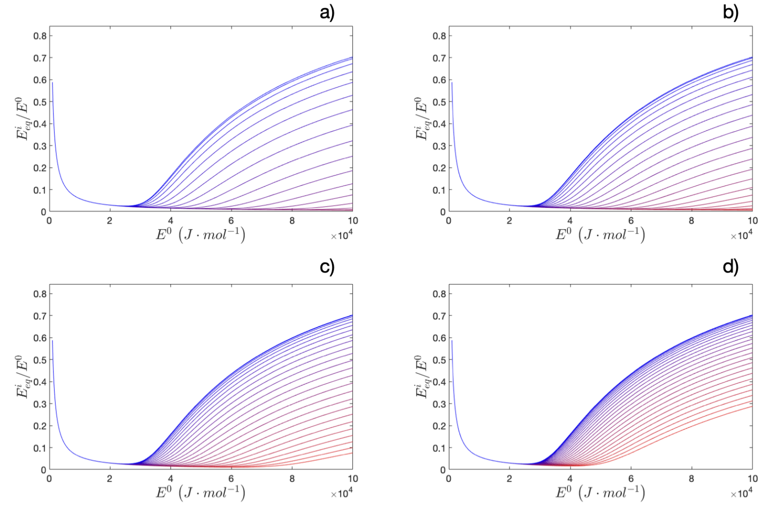

Figure S5: Available energy at population equilibrium ($E^i_{eq}/E^0$) as a function of $E^0$ for $N = 50$ consumers and $E^0_i$ distributions as in Fig. S2b for several widths: a) $\sigma = 0.5$ ; b) $\sigma = 1$ ; c) $\sigma = 2$ ; d) $\sigma = 4$. The remaining model parameters are common to all specialists. Due to symmetry, only species labeled 1-25 are shown. The color gradient indicates the species. Red line corresponds to label 1 and blue line corresponds to label 25.

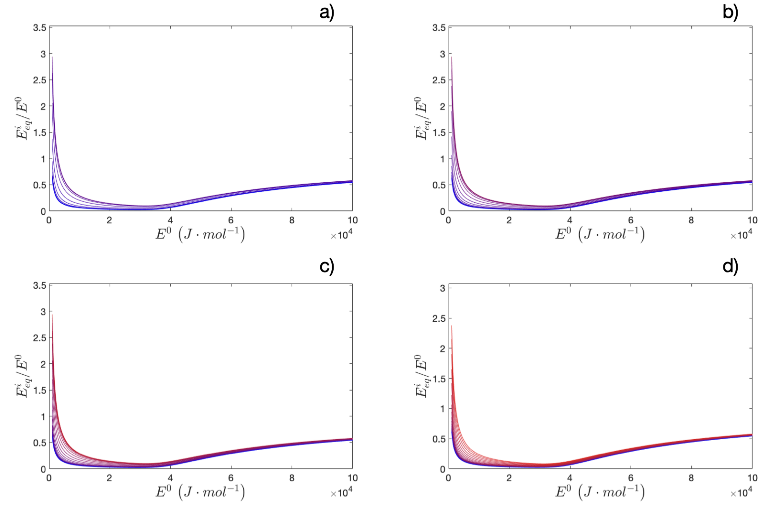

Figure S6: Available energy at population equilibrium ($E^i_{eq}/E_0$) as a function of $E^0$ for $N = 50$ consumers and $r_i$, $P_i$ distributions as in Fig. S2c and S2d for several widths: a) $\sigma = 0.5$ ; b) $\sigma = 1$ ; c) $\sigma = 2$ ; d) $\sigma = 4$. The remaining model parameters are common to all consumers. The color code as in Fig. S5.

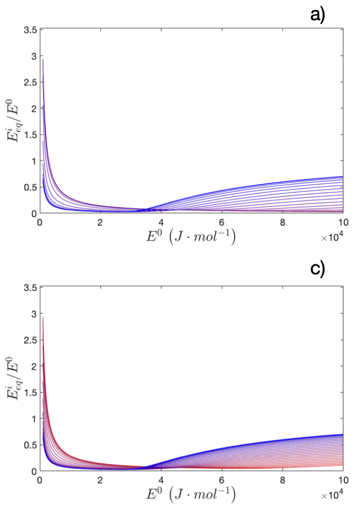

Figure S7: Available energy at population equilibrium ($E^i_{eq}/E^0$) as a function of $E^0$ for $N = 50$ consumers and for $\phi_i$, $E^0_i$, $r_i$ and $P_i$ distributions as in Fig. S2 for several widths: a) $\sigma = 0.5$ ; b) $\sigma = 1$ ; c) $\sigma = 2$ ; d) $\sigma = 4$. The color code as in Fig. S5.

#### 3.4 Relation between biological and power supply diversities

We consider the dependence of the biological diversity ($H_B$) on the power supply diversity ($H_P$). The values for both $H_B$ and $H_P$ are computed for energy scales ($E^0$) on the range $10^3 - 10^5\, J \cdot mol^{-1}$ and number of consumers ($N$) taking values on the range $10 - 1000$.

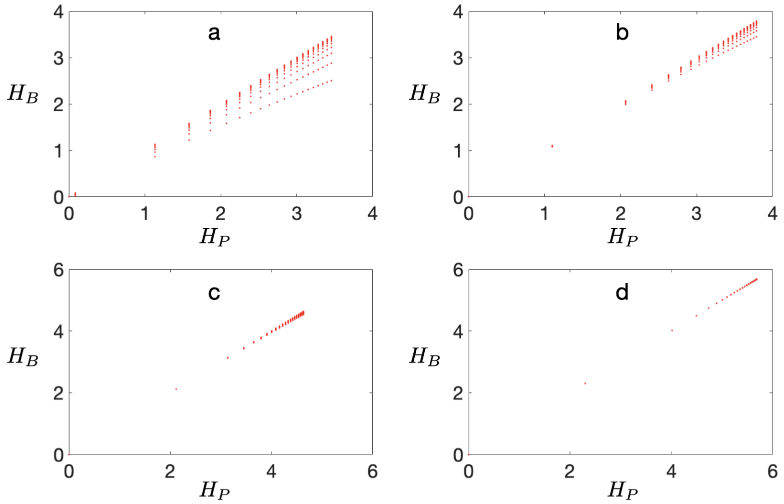

Figure S8: $H_B$ vs $H_P$ for a distribution of $\phi_i$ as in Fig. S2a and widths: a) $\sigma = 0.01$ ; b) $\sigma = 0.1$ ; c) $\sigma = 1$ ; d) $\sigma = 10$. The remaining model parameters are common to all consumers.

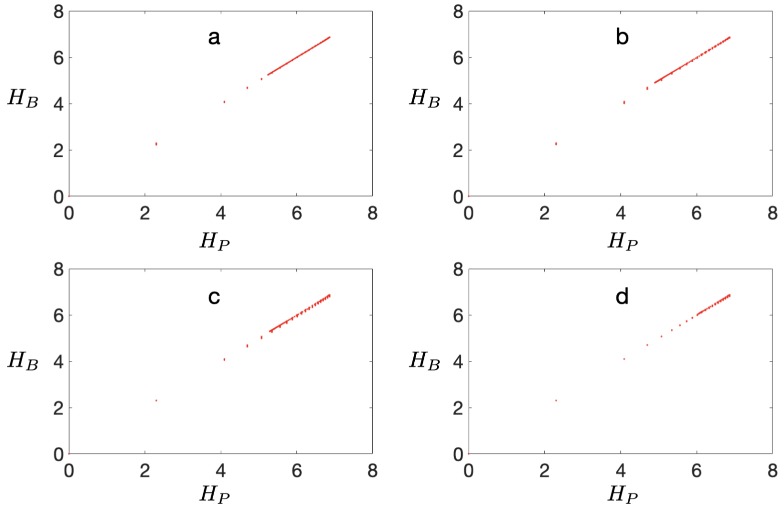

Figure S9: $H_B$ vs $H_P$ for a distribution of $E_i^0$ as in Fig. S2b and widths: a) $\sigma = 0.01$ ; b) $\sigma = 0.1$ ; c) $\sigma = 1$ ; d) $\sigma = 10$. The remaining model parameters are common to all consumers.

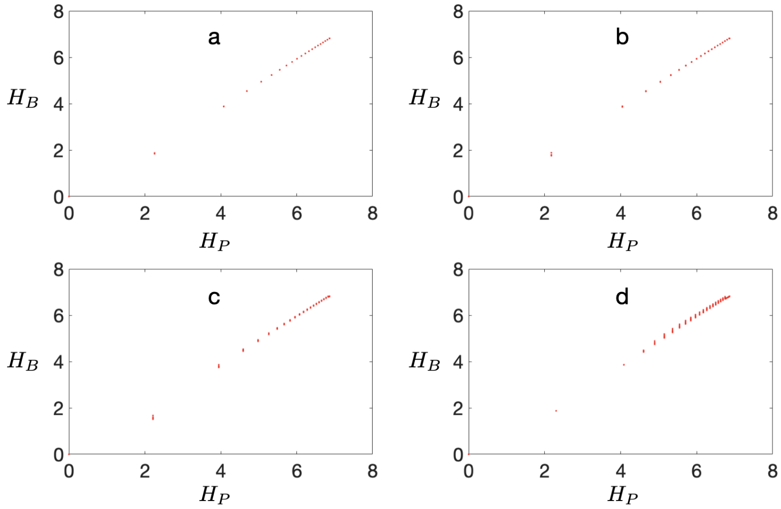

Figure S10: $H_B$ vs $H_P$ for distributions of uptake rate ($r_i$) and maintenance power ($P_i$) as in Fig. S2c and S2d and widths: a) $\sigma = 0.01$ ; b) $\sigma = 0.1$ ; c) $\sigma = 1$ ; d) $\sigma = 10$. The remaining model parameters are common to all consumers.

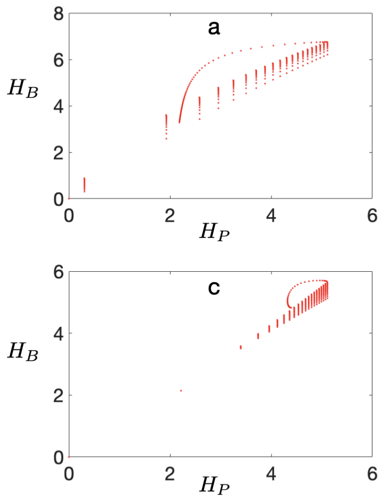

Figure S11: $H_B$ vs $H_P$ for distributions of $\phi_i$, $E_i^0$, $r_i$ and $P_i$ as in Fig. S2 and widths: a) $\sigma = 0.01$ ; b) $\sigma = 0.1$ ; c) $\sigma = 1$ ; d) $\sigma = 10$.

#### 3.5 Biological diversity dependence on pH

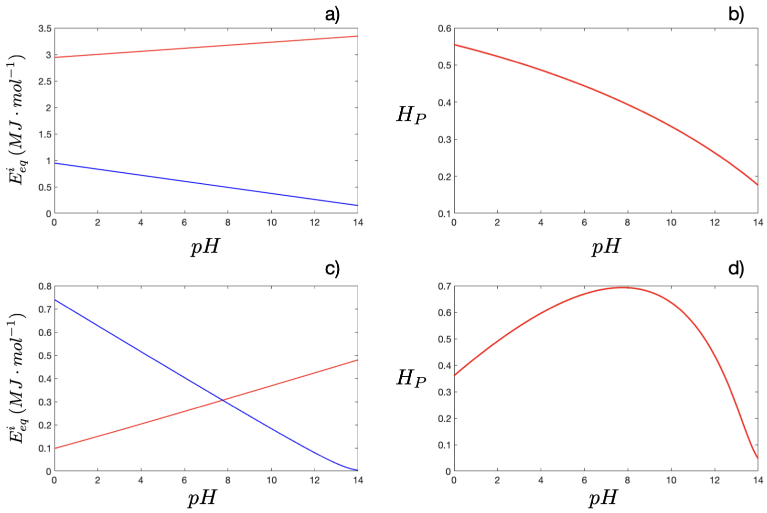

Figure S12: Available energy at equilibrium and power supply diversity as a function of the $pH$ for just 2 specialists, one being a consumer of $H^+$ and the other a producer of $H^+$. The temperature is set to $T = 300\,K$ and the stoichiometric coefficients are 5 (for the producer) and 10 (for the consumer). We consider a trade-off between uptake and power maintenance as in Fig. S2c andS2d, while all substrates have the same input concentration given by $\phi_0 = 5\frac{\frac{kP_0}{r_{max}RT}}{W\left(\frac{kP_0}{r_{max}RT}\, e^{\frac{r_{max}E^0 - P_0}{r_{max}RT}}\right)}$, where $W(z)$ is the Lambert $W$-function. The graphics show: a) and c) available energy at equilibrium for the $H^+$-consumer (blue line) and $H^+$-producer (red line) as a function of the $pH$ ; b) and d) show the corresponding power supply diversity.

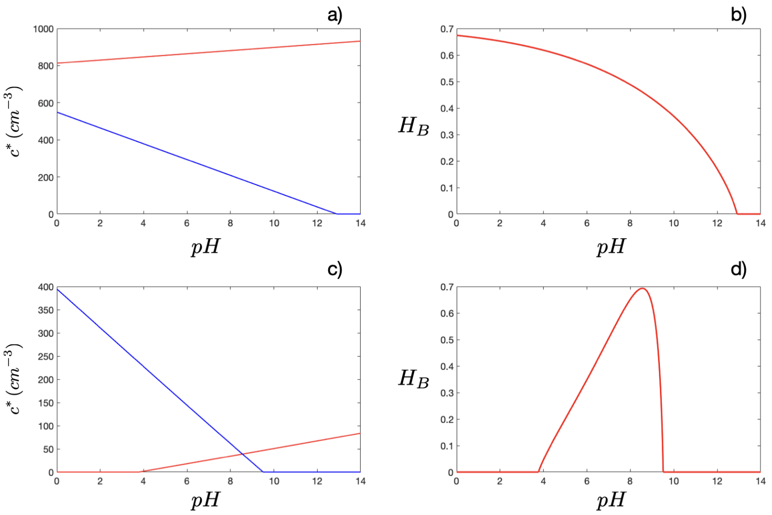

Figure S13: Cell abundance and biological diversity as a function of the pH corresponding to the data shown in Fig. S12. The graphics show: a) and c) stationary concentration of $H^+$-consumer (blue line) and $H^+$-producer (red line) as a function of the $pH$. The data shown in a) (resp. c)) corresponds to the available energy shown in Fig. S12a (resp. S12c); b) and d) show the corresponding biological diversity.
